# Adjacent terrestrial landscapes impact the biogeographical pattern of soil *Escherichia coli* in produce fields by modifying the importance of environmental selection and dispersal

**DOI:** 10.1101/2020.06.30.181495

**Authors:** Jingqiu Liao, Peter Bergholz, Martin Wiedmann

## Abstract

High-quality habitats for wildlife (e.g., forest) provide essential ecosystem services while increasing species diversity and habitat connectivity. Unfortunately, presence of such habitats adjacent to produce fields may increase risk for contamination of fruits and vegetables by enteric bacteria, including *Escherichia coli. E. coli* survives in extra-host environments (e.g., soil) and could disperse across landscapes by wildlife. Understanding how terrestrial landscapes impact the distribution of soil *E. coli* is of importance in assessing the contamination risk of agricultural products. Here, using multi-locus sequence typing, we characterized 938 *E. coli* soil isolates collected from two watersheds with different landscape patterns in New York state, USA, and compared the distribution of *E. coli* and the influence of two ecological forces (environmental selection and dispersal) on the distribution between these two watersheds. Results showed that for the watershed with widespread produce fields, sparse forests, and limited interaction between the two land-use types, *E. coli* composition was significantly different between produce field sites and forest sites; this distribution was shaped by relatively strong environmental selection likely from soil phosphorus and slight dispersal limitation. For the watershed with more forested areas and stronger interaction between produce field sites and forest sites, *E. coli* composition between these two land-use types was relatively homogeneous; this distribution appeared to a consequence of wildlife-driven dispersal, inferred by competing models. Collectively, our results suggest that terrestrial landscape attributes could impact the biogeographic pattern of enteric bacteria by adjusting the importance of environmental selection and dispersal.

**IMPORTANCE:** Understanding the ecology of enteric bacteria in extra-host environments is important to allow for development and implementation of strategies to minimize pre-harvest contamination of produce with enteric pathogens. Our findings suggest that watershed landscape is an important factor influencing the importance of ecological drivers and dispersal patterns of *E. coli*. For watersheds with widespread produce fields, *E. coli* appears to experience local adaptation, possibly due to exposure to environmental stresses associated with agricultural activities. In contrast, for watersheds with high forest coverage we found evidence for wildlife-driven dispersal of *E. coli*, which might facilitate more frequent genetic exchange in this environment. Agricultural areas in such watersheds may have a higher risk of produce contamination due to less environmental constraints and higher potential of dispersal of enteric bacteria between locations. The significance of our research lies in exploring ecological principles underlying the biogeographic pattern of enteric bacteria at the regional level, which can inform agricultural, environmental and public health scientists that aim to reduce the risk of food contamination by enteric bacteria.

## INTRODUCTION

Forests and other riparian buffers can provide ecological and agricultural benefits (e.g., reducing soil erosion and leaching of chemical and fecal waste into surface water sources, providing habitat and connective pathways for wildlife) as well as aesthetic benefits (1–3). While high quality habitats (e.g., forest) offer conservation services, they may also bring unintended consequences and may increase the risk of pre-harvest contamination of produce crops. For one, extra-host environments, such as soil in high quality habitats, can be a critical reservoir for enteric bacteria, leading to potential dispersal of enteric bacteria to adjacent agricultural fields (4). Direct fecal deposition onto produce by wildlife is also a potential pathway introducing enteric bacteria onto food crops (5). By providing wildlife movement pathways, high quality habitats may facilitate the wildlife-driven dispersal of enteric bacteria through riparian corridors to agricultural regions, possibly resulting in contamination of food crops (6–9). Subsequent persistence and/or regrowth of pathogenic enteric bacteria introduced into fresh produce fields would then further increase food safety risks (10). Since the survival of enteric bacteria and movement of wildlife vary by land use types (11, 12), it is reasonable to hypothesize that watershed landscape impacts the distribution of enteric bacteria.

Environmental selection and dispersal are two fundamental ecological forces that drive the distribution of bacteria (13–16). The essential roles of environmental selection via abiotic (e.g., pH, salinity) and biotic selective pressures on bacteria have been well documented in many local and even global habitats (17–20). Environmental selection facilitates the genetic divergence of some ecophysiological traits owing to their contribution to fitness benefits for adaptation of bacteria to diverse habitats, such as those with different land cover types (21, 22). With the influence of environmental selection, a high level of bacterial dissimilarity between locations (beta diversity) can be maintained in a wide range of environments (15, 23, 24).The role of dispersal in driving the distribution of bacteria at local as well as regional scales is evident since dispersal provides a mechanism for bacteria to colonize new habitats (25, 26). The relative importance of dispersal in shaping bacterial distribution varies among microbial taxa due to diversity in the capacity of bacteria to disperse via wind, water, and wildlife. For example, bacteria with a long range dispersal capacity (e.g., *Polaromonas*) tend to exhibit a more global distribution (27), while bacteria with a limited dispersal range (e.g., *Rhizobiaceae, Bradyrhizobiaceae, Xanthomonadaceae*) tend to show more ecological specialization (15). Importantly, wildlife presence and movement is fundamentally affected by the physical elements and features of land (15). Thus, wildlife-driven dispersal of bacteria can be quantitatively predicted by landscape ecological methods based on the relationship of wildlife behaviors and landscape characteristics (e.g., patchiness, land-use interspersion, patch connectivity, patch diversity, and land-use interactions) (8, 28–30). Based on these principles, investigating environmental selection and dispersal of enteric bacteria from and within habitats with distinct landscape patterns has the power to elucidate the role of terrestrial landscapes in impacting the distribution of enteric bacteria and assess the associated risk of preharvest contamination of food by pathogenic enteric bacteria.

As a commensal or pathogenic enteric bacterium widespread in diverse habitats, *Escherichia coli* primarily resides in the intestines of warm-blooded animals, and survives in extra-host environments such as water, soil, sediments as well (31, 32). Soil is a habitat of particular interest for *E. coli*, since the high chemical and physical heterogeneity of soil across different environments could pose multifarious environmental selection pressures on *E. coli* (22, 23, 32). Observations that the prevalence of *E. coli* varies by land cover types (e.g., deciduous forest, cropland, pasture) (22) also suggest that different land uses could stimulate different types and intensities of selective pressures that act on *E. coli*. The key edaphic variables influencing the growth of *E. coli* in soil are commonly recognized as pH and moisture (23, 33), while some other soil properties such as organic matter and texture could also play a role (10). In addition, wildlife, such as avian species and ruminant animals, could act as dispersal vehicles of *E. coli* (34, 35). *E. coli* can also be transmitted between wildlife hosts through contact and can be deposited in new locations (e.g., produce fields) by defecation, which often happens when wildlife forages for food (36). Given the intensive interaction with both extra-host and host habitats, *E. coli* may be a useful model to build predictive capabilities surrounding interactions between bacteria and agricultural landscapes at the meter to kilometer scale. Such an understanding is particularly important for lands where fresh fruits and vegetables are cultivated, as it could be used to develop better strategies for minimizing pathogen introduction into preharvest environments.

We hypothesized that the importance of environmental selection and dispersal for the distribution of *E. coli* is dependent on landscape and specifically hypothesized that (i) *E. coli* in watersheds with higher coverage of agricultural environments is strongly driven by environmental selection associated with agricultural activity, while (ii) *E. coli* in regions with higher coverage of natural environment is largely influenced by wildlife-driven dispersal. To test our hypotheses, we characterized 938 generic *E. coli* isolates obtained from soil samples collected from two watersheds - Flint Creek and Hoosic River, both located in the New York State, using a hierarchical multi-locus sequence typing (MLST) scheme. These two watersheds represent an interesting comparison between one with widespread produce fields and limited interaction between produce fields and forest (Flint Creek; 69% produce field, 12% forest by area; adjacency_produce|forest_ = 23%; Fig. 1a) and one with heavily forested areas and strong interaction between produce fields and forest (Hoosic River; 28% produce field, 38% forest by area; adjacency_produce|forest_ = 36%; Fig. 1b). Next, we investigated the distribution of *E. coli* in these two watersheds, and assessed the relationship between *E. coli* distribution and soil variables and the distance-decay relationship among *E. coli* populations. Last, we developed dispersal models for four wildlife vehicle candidates (large nuisance wildlife species, small mammals, small flocking insectivore/granivores, and migratory bird flocks) to quantify the importance of wildlife-driven dispersal on the dispersal of *E. coli* in the two watersheds.

**FIG 1.**
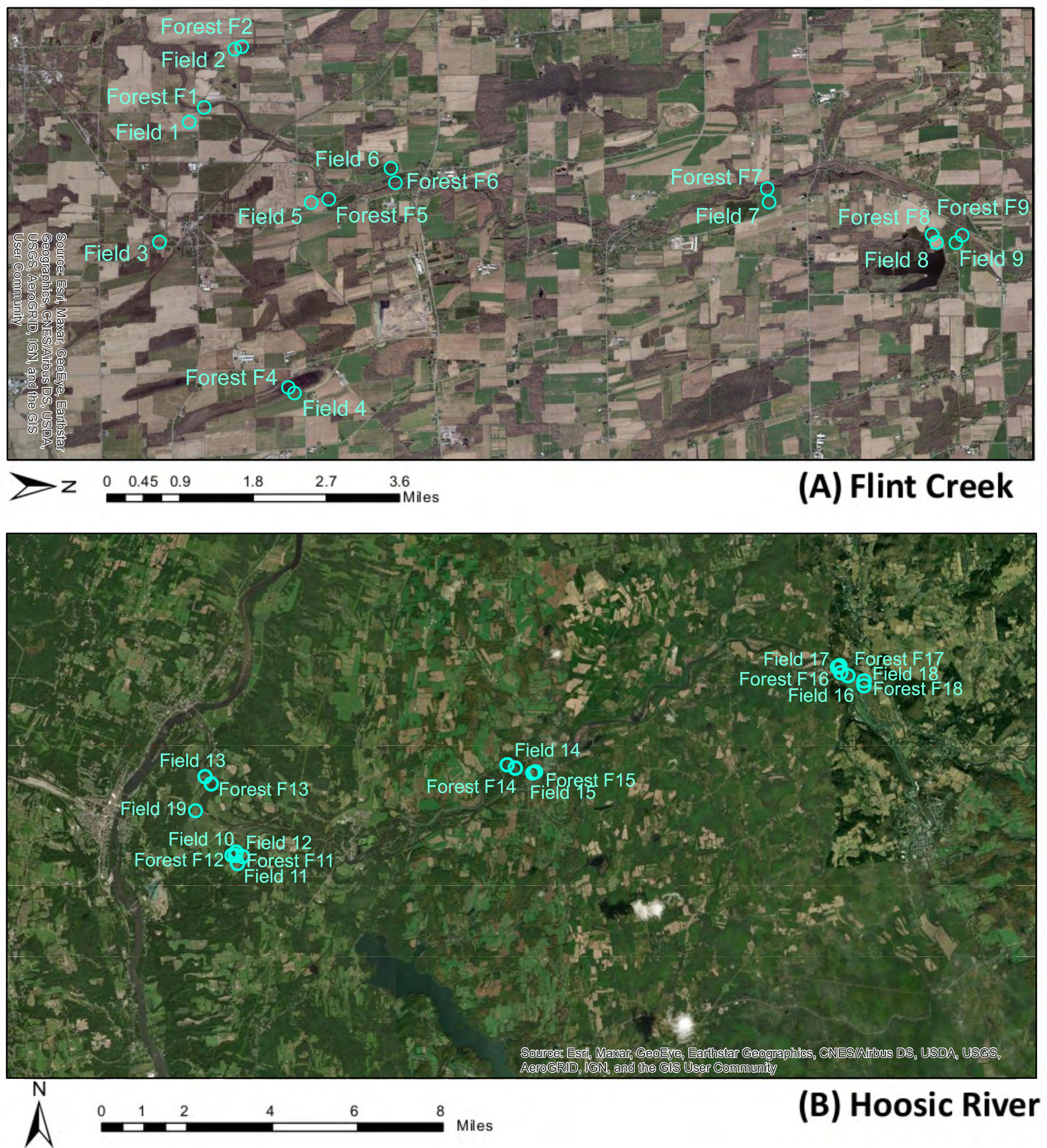
Sampling maps of (a) Flint Creek and (b) Hoosic River. Dots indicate the sampling sites within each watershed. Map layers for land cover (National Land Cover Database [NLCD], 2006) were acquired from the U.S. Geological Survey (USGS) Earth Explorer geographical data bank (http://earthexplorer.usgs.gov/).

## RESULTS

### Distribution of *E. coli*

Soil samples from Flint Creek (predominated by produce fields) showed considerably lower prevalence of *E. coli* than soil samples from Hoosic River (predominated by forests); 35% and 72% of soil samples, respectively, were positive for *E. coli* in these two watersheds. These samples yielded 289 and 649 *E. coli* isolates, respectively. Based on initial 2-gene MLST (*mdh* and *uidA*), a total of 138 isolates from Flint Creek and 277 isolates from Hoosic River were selected for characterization by full 7-gene MLST (*aspC, clpX, icd, lysP, fadD*, in addition to *mdh* and *uidA*). The 7-gene MLST generated 121 unique multilocus sequence types (ST) for Flint Creek and 191 unique ST for Hoosic River (Table S1, Table S2). Analysis by goeBURST identified 96 and 108 *E. coli* clonal groups for Flint Creek and Hoosic River, respectively, based on ST at single locus variant level (Fig. S1, Fig. S2). For Flint Creek, forest sites had slightly higher mean richness of *E. coli* clonal groups than produce field sites, but the difference was not significant (*p* = 0.86) (Fig. S3). In contrast, the mean richness of *E. coli* clonal groups from produce field sites was significantly higher than forest sites in the Hoosic River watershed (*p* < 0.05) (Fig. S3).

*E. coli* in the two watersheds displayed distinct distribution patterns. For Flint Creek, principal coordinates analysis (PCoA) based on the dissimilarity of *E. coli* clonal groups clustered sampling sites by land-use (Fig. 2a). Both permutational multivariate analysis of variance (PERMANOVA) test and analysis of similarities (ANOSIM) test showed that this clustering by land-use was significant (*p* < 0.05; Table S3, Table S4). By contrast, for the Hoosic River, the sampling sites were not significantly clustered by land-use in PCoA (Fig. 2b) of *E. coli* clonal groups (PERMANOVA *p* = 0.86, ANOSIM *p* = 0.58; Table S3, Table S4), indicating a more homogeneous composition of *E. coli* between produce field sites and forest sites in this watershed. To test the robustness of these results and to assess the effects of potential sampling bias on the *E. coli* diversity, we further repeated PCoA, PERMANOVA, and ANOSIM analyses after excluding sites with a low number of *E. coli* clonal groups detected (≤ 3); sites being excluded were Field 6, Field 8, and Forest F9 from Flint Creek. Results showed that sampling sites from this subset were significantly clustered by land-use in the PCoA plot (Fig. S4) for *E. coli* clonal groups from Flint Creek at least at the 0.1 level (PERMANOVA *p* = 0.05, ANOSIM *p* = 0.08; Table S5). Slightly larger *p* values in PERMANOVA and ANOSIM tests using a subset of sites (as comparted to *p*-values for the whole set of sites) are likely due to the reduced sample size in these analyses.

**FIG 2.**
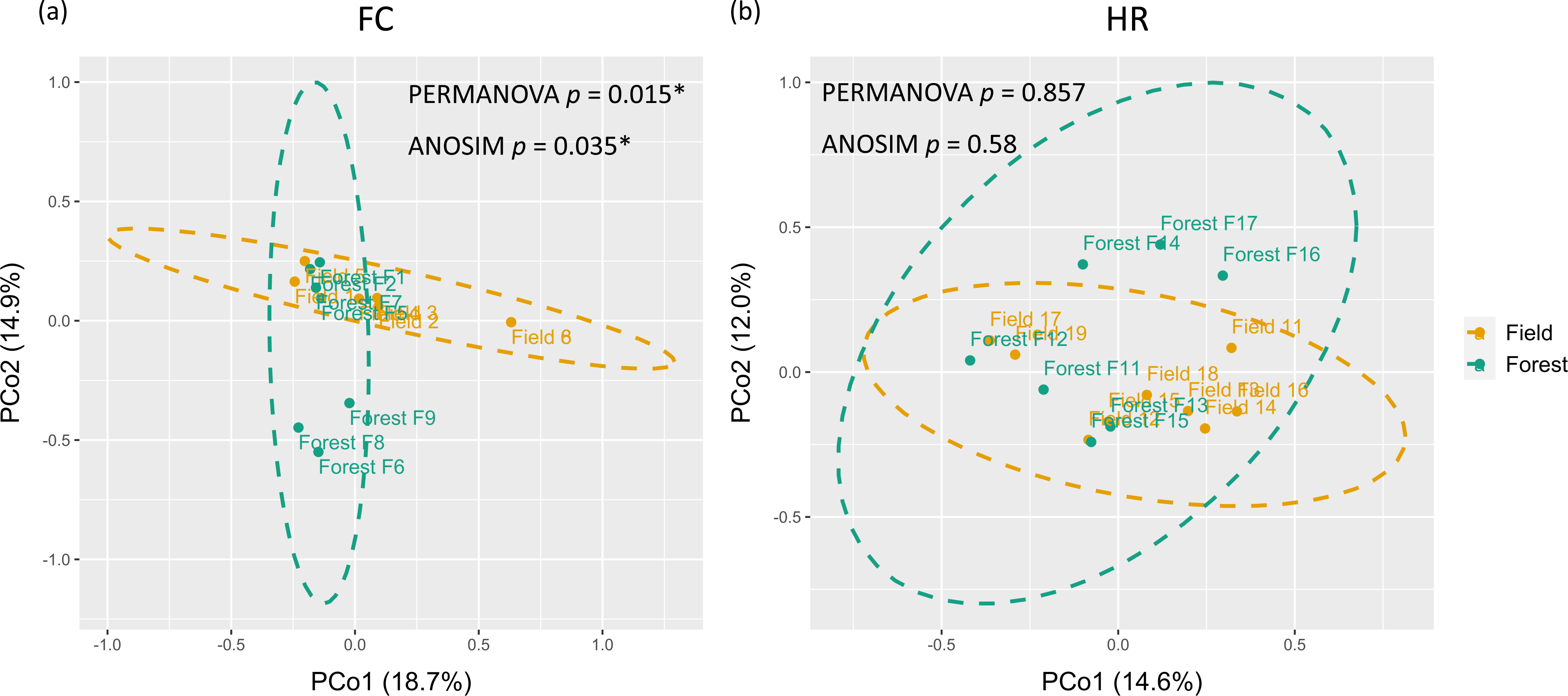
PCoA plots of *E. coli* clonal groups of sites in (a) Flint Creek (FC) and (b) Hoosic River (HR). Green dots indicate forest sites; orange dots indicate produce field sites; Green circle indicates the 95% confidence ellipse of forest sites; orange circle indicatedsthe 95% confidence ellipse of produce field sites. *p* values of permutational multivariate analysis of variance (PERMANOVA) test and analysis of similarities (ANOSIM) test are shown; “*” indicates the clustering of sampling sites by land-use is significant at the 0.05 level. (a) PCo Axis 1 and 2 explained 18.7% and 14.9%, respectively, of the variation of *E. coli* clonal groups for Flint Creek. (b) PCo Axis 1 and 2 explained 14.6% and 12.0%, respectively, of the variation of *E. coli* clonal groups for Hoosic River.

### The importance of soil variables driving the distribution of *E. coli*

We first employed variance partitioning analysis (VPA) to quantify the contribution of soil variables and spatial variables to the distribution of *E. coli* from Flint Creek and Hoosic River. After screening for high levels of covariation, moisture, pH, sodium, phosphorus, barium, manganese, and antimony were included in VPA (Table S6). VPA revealed that the soil variables individually explained 11.7% of the biological variation of dissimilarity of *E. coli* clonal groups from Flint Creek, while spatial variables individually explained 5.5% of this variation (Fig. 3a). The variation explained by spatially structured soil variables was relatively low (0.1%) and about 83% of the variation was unexplained by selected variables (Fig. 3a). By contrast, for *E. coli* clonal groups from Hoosic River, about 99% of the variation of dissimilarity could not be explained by selected variables (Fig. 3b). Individually, selected soil variables did not explain any of the observed variation and spatial variables only explained 1% of the variation for the Hoosic River watershed (Fig. 3b). These results indicate that environmental selection outweighs spatial factors with regard to the importance for *E. coli* distribution in the Flint Creek watershed; *E. coli* from this watershed appear to undergo much stronger environmental selection or a much slower rate of dispersal than *E. coli* in the Hoosic River watershed.

**FIG 3.**
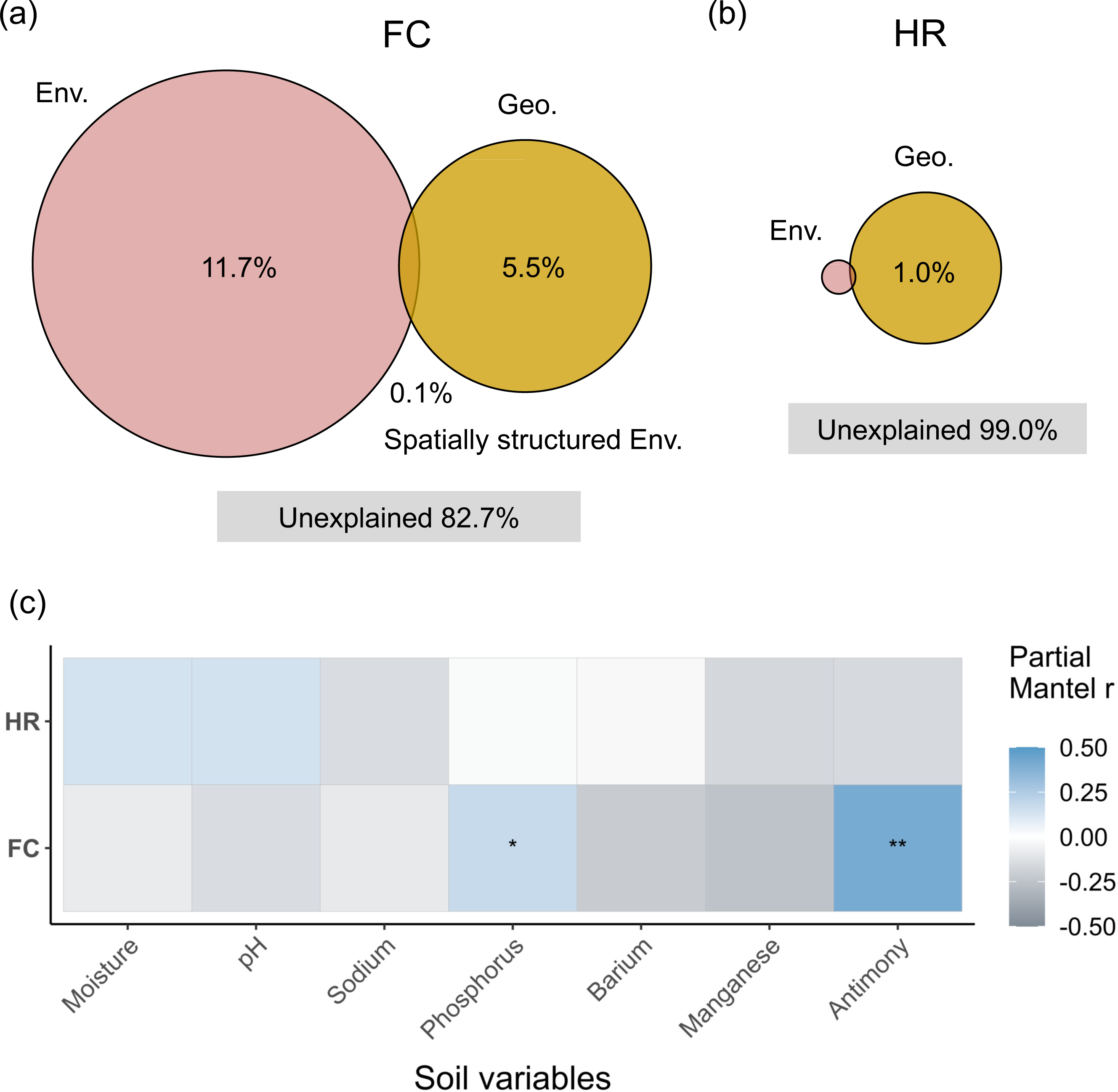
Variance partitioning analysis (VPA) showing relative contributions of spatial factors (Geo.), soil variables (Env.), and spatially structured environmental variables to the variations of the dissimilarity of *E. coli* clonal groups based on Bray-Curtis distance for (a) Flint Creek (FC) and (b) Hoosic River (HR). (c) Partial Mantel correlation between the dissimilarity of *E. coli* clonal groups and geographic distance-correlated dissimilarity of soil variables for FC and HR. r is the Partial Mantel correlation coefficient. Significant correlations at the 0.05 and 0.01 level are denoted by “ *” and “ * *”, respectively.

To identify the key soil variables correlated with the dissimilarity of *E. coli* clonal groups, we performed partial Mantel tests by correcting the correlation of geographic distance and biological dissimilarity. For Flint Creek, partial Mantel tests showed that geographic distance-corrected phosphorus and antimony concentrations in soil were significantly correlated with the dissimilarity of *E. coli* clonal groups (correlation coefficient r = 0.18, *p* < 0.05 and r = 0.41, *p* < 0.05, respectively) (Fig. 3c). In addition, for sites in Flint Creek, we observed that the mean concentration of phosphorus was significantly higher for soil samples from produce field sites as compared to samples from forest sites (Mann-Whitney test *p* < 0.05, Fig. S5a). Also, in Flint Creek, the mean concentration of antimony was higher for soil samples from produce field sites compared to samples from forest sites, but the difference was not significant (Mann-Whitney test *p* = 0.45, Fig. S5b). In contrast, none of the soil variables were found to be significantly correlated with the dissimilarity of *E. coli* clonal groups from Hoosic River (*p* > 0.05) (Fig. 3c). This result was consistent with our observation, based on VPA, that soil variables minimally contributed to the distribution of *E. coli* clonal groups in the Hoosic River watershed.

### The relationship between the dissimilarity of *E. coli* and geographic distance

We first assessed the relationship between dissimilarity of *E. coli* clonal groups and geographic distance using Mantel tests. Results showed a weak correlation between the dissimilarity of *E. coli* clonal groups and geographic distance for Flint Creek at the 0.1 level (Mantel correlation coefficient r = 0.16, *p* = 0.08; Table S7). By contrast, no significant correlation at the 0.1 level was observed between the dissimilarity of *E. coli* clonal groups and geographic distance for Hoosic River (r = 0.11, *p* = 0.12; Table S7). Linear regression analysis further showed that the slope of the regression line of the linear relationship between the dissimilarity of *E. coli* clonal groups and geographic distance for Flint Creek (slope = 3.4 × 10^-3^, R^2^ = 0.027, Fig. 4a) was about 3 times steeper than that for Hoosic River (slope = 9.7 × 10^-4^, R^2^ = 0.011, Fig. 4b), indicating a stronger distance-decay relationship in *E. coli* in Flint Creek than Hoosic River. Overall, results of Mantel test and linear regression analysis along with the VPA suggest that spatial factors play a more important role in driving the distribution of *E. coli* clonal groups in Flint Creek than Hoosic River, and the dispersal of *E. coli* in Flint Creek was slightly limited, while *E. coli* in the Hoosic River was more likely not constrained by dispersal limitation.

**FIG 4.**
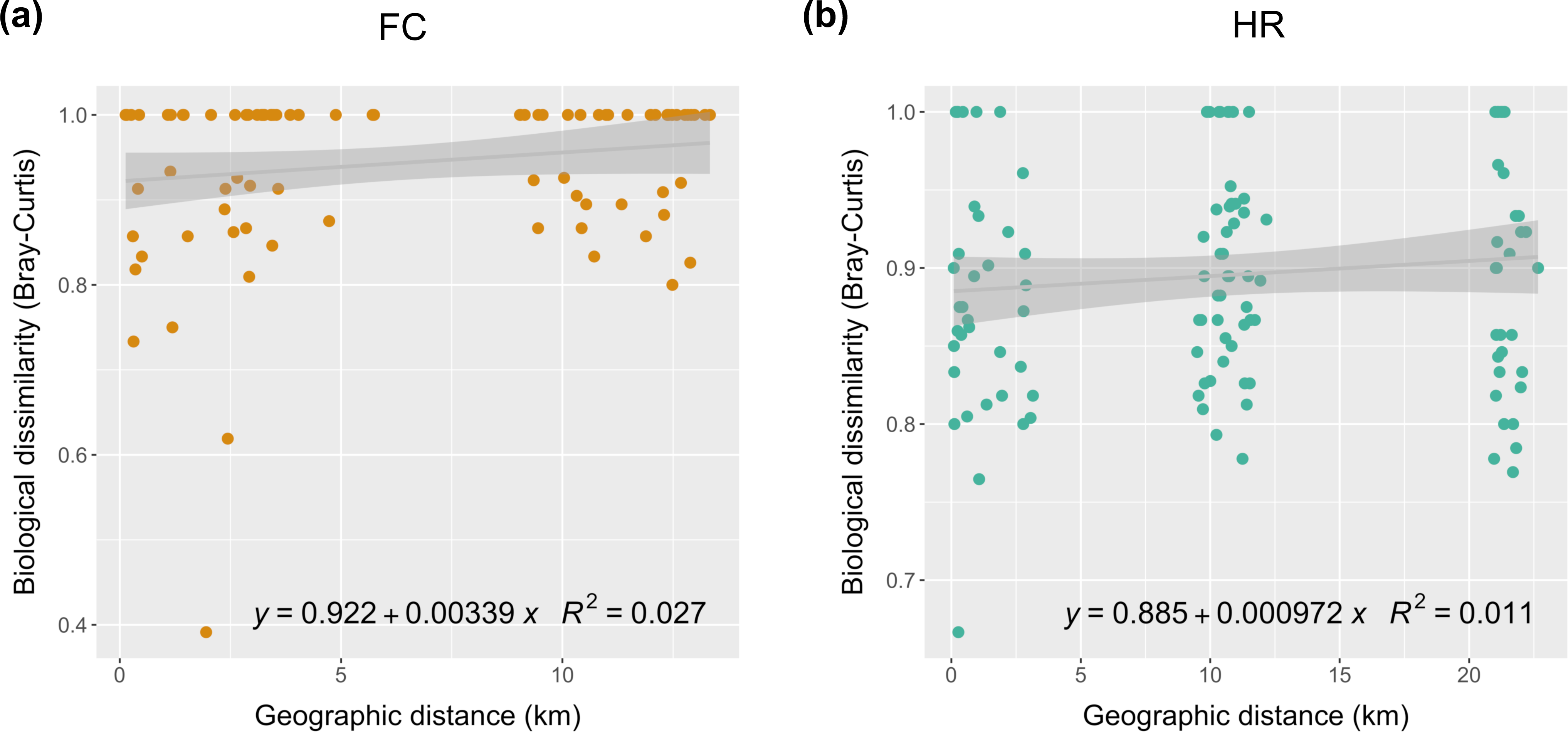
Linear relationship between biological dissimilarity of *E. coli* clonal groups and geographical distance for (a) Flint Creek (FC) and (b) Hoosic River (HR). The biological dissimilarity of *E. coli* clonal groups was calculated as Bray–Curtis distance. Geographical distance was calculated as the actual physical distance. Linear regression line is in grey; shaded area indicates 95% confidence region; R^2^ indicates the variability explained by fitted linear regression model and the formula of the linear relationship is shown.

To evaluate the impact of land-use on the relationship between *E. coli* and geographic distance, we conducted Mantel tests and assessed the distance-decay relationship for produce field sites and forest sites separately within each watershed. For Flint Creek, Mantel tests showed that the correlation between the dissimilarity of *E. coli* clonal groups and geographic distance was not significant at the 0.1 level; the correlation coefficient for forest sites (r = 0.27) was slightly larger than that for produce field sites (r = 0.22) (Table S8). For Hoosic River, the dissimilarity of *E. coli* clonal groups and geographic distance for forest sites was significantly and highly correlated (*p* < 0.05, correlation coefficient = 0.49), while there was no significant correlation for produce field sites (*p* = 0.56) (Table S8). Consistently, based on the linear regression analysis assessing the distance-decay relationship, the slope of the linear regression line for forest sites in both Flint Creek and Hoosic River (slope = 7.0 × 10^-3^ and 4.3 × 10^-3^, respectively) was steeper than that for produce field sites (slope = 5.1 × 10^-3^ and -3.0 × 10^-4^, respectively) (Fig. S6a and S6b). These results suggest that dispersal limitation for *E. coli* tend to be weaker in produce field sites than in forest sites, which implies that dispersal of *E. coli* may be more efficient in the produce fields.

### Wildlife-driven dispersal of *E. coli*

Four common classes of wildlife vehicles (large nuisance wildlife species, small mammals, small flocking insectivore/granivores, and migratory bird flocks) were selected for identifying potential dispersal vehicles of *E. coli* (characteristics of these dispersal vehicles are detailed in Table 1). By adjusting distances among sampled sites to account for movement preferences of these four types of wildlife vehicles (*i.e.*, cost-distance or landscape resistance modeling), we sought to assess whether dispersal associated with wildlife behavior explains the *E. coli* distribution better than distance alone. As shown in Table 2, the predicted dispersal model was developed based on the most-likely cost and attraction models selected for each wildlife vehicle according to their characteristics. The predicted dispersal model for small mammals was defined to have a biological riparian corridor effect, no proximity effect, absolute dispersal barriers effect, and no attraction coefficient. The predicted dispersal model for large nuisance wildlife species was defined to have a biological riparian corridor effect, strong proximity effect, porous dispersal barriers effect, and habitat quality coefficient. The predicted dispersal model for migratory bird flocks was defined to have a biological riparian corridor effect, weak proximity effect, no dispersal barriers effect, and area independent coefficient. The predicted dispersal model for small flocking insectivore/granivores was defined to have a biological riparian corridor effect, weak proximity effect, absolute dispersal barriers effect, and area independent coefficient. Definition of these effects can be found in Table 1.

**TABLE 1.** Basic dispersal model characteristics

| Class | Description |
| --- | --- |
| **Dispersal vehicles<sup>a</sup>** |  |
| LT: Large nuisance wildlife species (e.g. "deer" or "feral swine") | Forests, scrublands, grasslands, wooded wetlands and cultivated croplands impose low movement costs, roads impose increased movement costs based on census category |
| ST: Small mammals (e.g. "shrews", "voles" and "mice") | Light urban development, grassland and scrublands impose low movement costs, roads impose increased movement costs based on census category |
| LB: Small flocking insectivore/granivores (e.g., "starlings") | Open water and heavy urban development are higher cost, all other movement costs are low |
| MB: Migratory bird flocks (e.g. "Canada goose") | Heavy urban development imposes higher cost, all other movement costs are low. |
| **Riparian corridor** (movement costs are reduced by half) |  |
| Adjacency | All land parcels that overlap a 100 m zone around the main river/creek |
| Distance | Land within 100 m of the main river/creek |
| Biological | Land below the 50 yr flood height for the main river/creek and adjacent wetlands |
| None | No riparian corridor effect |
| **Dispersal barriers** |  |
| Absolute | Major roads and waterways are absolute barriers (movement cost 40,000 per pixel) |
| Porous | Major roads and waterways are porous barriers (movement cost 200 per pixel) |
| None | No barrier effect |
| **Proximity effects** (specifics vary by vehicle) |  |
| Strong | Nearness to high quality habitat substantially reduces movement cost |
| Weak | Nearness to high quality habitat somewhat reduces movement cost |
| None | No benefit of proximity to good cover |
| **Attraction (gravity) coefficients** (specifics vary by vehicle) |  |
| Habitat Quality | Proximity, interspersed and area of high quality habitat increase the chances that *E. coli* will be deposited |
| Reduced Habitat Effect | Effect of high quality habitat is reduced by half |
| Area Independent | As habitat quality model, but area of high quality habitat does not impact the result |
| None | Attraction does not influence dispersal |
| **Load (source) coefficient** |  |
| *E. coli* load estimation | Areas close to forest or pasture-class landcover are higher load. |

<sup>a</sup>Conceptual models are based on published literature (80-91).

**TABLE 2.** The predicted dispersal model for each wildlife vechile

| Vehicles | Most-likely cost model<sup>a</sup> |  |  | Most-likely attraction model (coefficient)<sup>b</sup> |
| --- | --- | --- | --- | --- |
|  | Riparian corridor | Proximity effects | Dispersal barriers |  |
| ST (Small mammals) | Biological | None | Absolute | None |
| LT (Large nuisance wildlife species) | Biological | Strong | Porous | Habitat Quality |
| MB (Migratory bird flocks) | Biological | Weak | None | Area Independent |
| LB (Small flocking insectivore/granivores) | Biological | Weak | Absolute | Area Independent |

<sup>a</sup>Cost model was developed based on three factors, riparian corridor, proximity effects, and dispersal barriers. Definition of the three factors can be found in Table 1. The most-likely cost model was selected based on the characteristics of each wildlife vehicle (Table 1).
<sup>b</sup>Definition of classes of attraction model can be found in Table 1. The most-likely attraction model was selected based on the characteristics of each wildlife vehicle (Table 1).

Mantel tests showed that none of these dispersal models significantly predicted the composition of *E. coli* clonal groups in Flint Creek (*p* > 0.05), while two wildlife-driven dispersal models – dispersal via migratory bird flocks (M459) and via small flocking insectivore/granivores (M556) – were found to be significantly correlated with the dissimilarity of *E. coli* clonal groups in Hoosic River (correlation coefficient r = 0.17 and r = 0.16, respectively; *p* < 0.05) (Fig. 5). In addition, the model for dispersal via large nuisance wildlife species (M377) was marginally significantly correlated with the dissimilarity of *E. coli* clonal groups (r = 0.14, *p* = 0.056), so the role of dispersal of *E. coli* by large nuisance wildlife species cannot be absolutely excluded. Thus, migratory bird flocks, small flocking insectivore/granivores, and large nuisance wildlife species were identified as potential dispersal vehicles which were associated with the distribution of *E. coli* in Hoosic River. The observation that cost-distance model correlated with the dissimilarity of *E. coli* clonal groups in Hoosic River better than geographic distance alone (Mantel correlation coefficient r = 0.11, *p* = 0.12; Table S7) suggests some dispersal among sites by the action of wildlife. Our results also suggest that wildlife-driven dispersal played a more important role in shaping the distribution of *E. coli* in Hoosic River as compared to Flint Creek.

**FIG 5.**
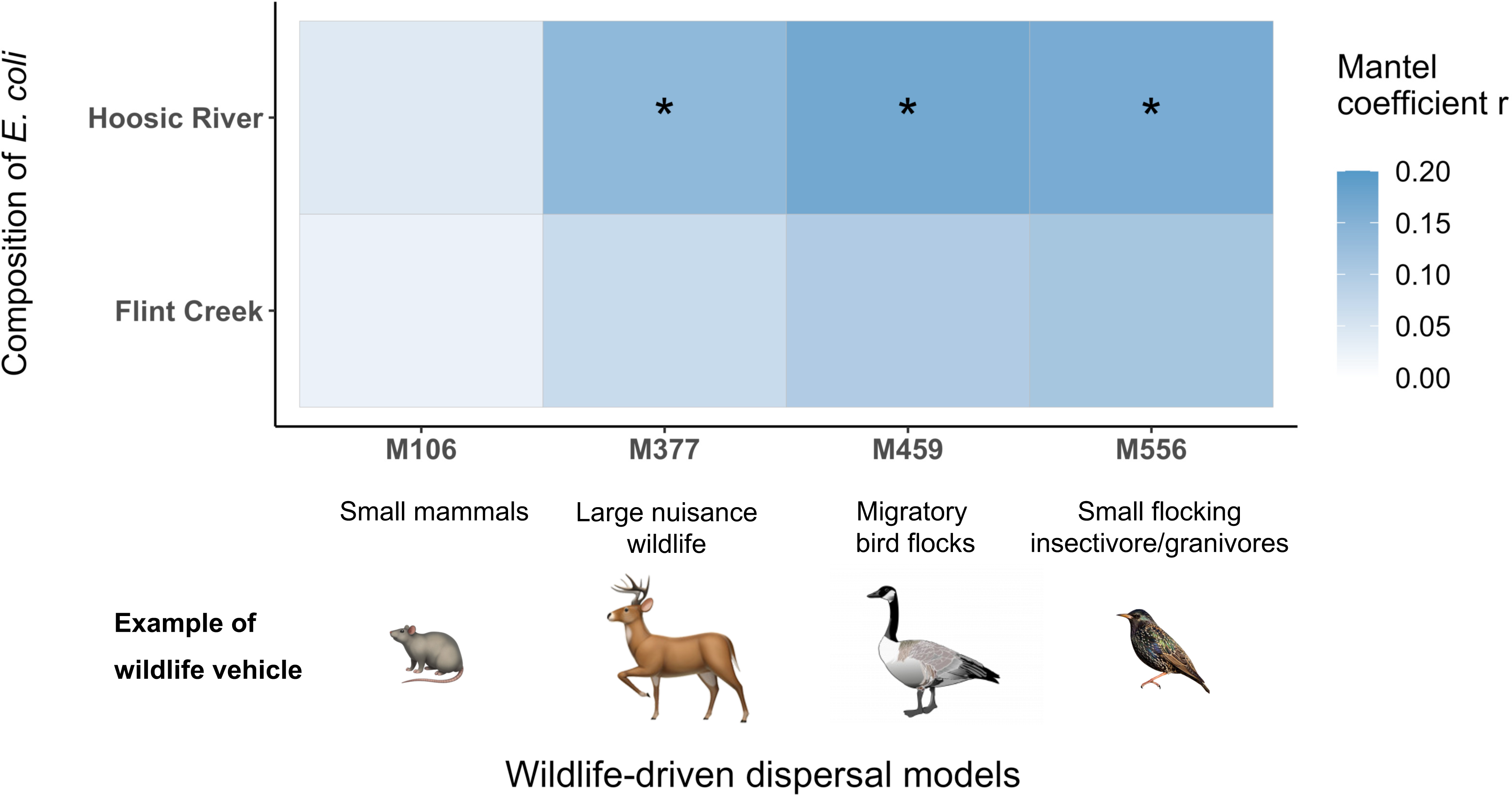
Mantel test result of wildlife-driven dispersal models and composition of *E. coli* clonal groups for Flint Creek and Hoosic River. M106, M377, M459, and M556 on the x-axis are the identification numbers of the predicted dispersal models for small mammals, large nuisance wildlife, migratory bird flocks, and small flocking insectivore/granivores, respectively. Examples of each wildlife vehicle type are shown. The description of the predicted dispersal model for each wildlife vehicle is detailed in Table 2. Models significant or marginally significant at the 0.05 level in Mantel tests are denoted by “ *”.

## DISCUSSION

*E. coli* has widely been used as an indicator of fecal contamination (37) and potential presence of other pathogenic enteric bacteria in water (38). *E. coli* comprises a wide spectrum of phenotypes including harmless commensal variants as well as distinct pathotypes with the capacity to either cause intestinal or extraintestinal infections in humans and many animals (39). The fecal-oral transmission route of *E. coli* often involves transient presence in extra-host habitats (e.g., surface water, soil, plant surfaces), including produce fields (23). Therefore, understanding the ecology of *E. coli* in extra-host habitats will not only provide an improved understanding of *E. coli* interaction with environment, but will also benefit public health by providing knowledge that can be used to minimizing introduction of *E. coli* and possibly other enteric pathogens into preharvest environments.

Environmental stressors such as limited availability of nutrients and water, presence of toxic molecules, and large alterations in temperature and moisture can impose fitness cost on *E. coli* and other microbes (40). Fragmented landscapes with smaller forest and grassland patches expose surface soil to sunlight and greatly increase daily variation in soil conditions. Reduced forest and grassland cover could also hinder the movement of wildlife, bringing negative demographic and genetic consequences (41). Thus, in order to disperse to and survive new habitats, *E. coli* needs to overcome those barriers by maintaining variable survival strategies such as evolving adaptive traits relying on dispersal to rescue local populations. In this scenario, landscape structure imposes constraints on environmental selection and dispersal, which is particularly essential for the dispersal of *E. coli* among different extra-host habitats.

To quantitatively probe the importance of environmental selection and dispersal in driving the distribution and composition of *E. coli* in soil under the impact of landscape, we compared the biogeographic patterns of *E. coli* isolated from two watersheds with distinct landscape patterns (i.e., Flint Creek, an area with widespread produce fields and limited interaction between produce fields and forest, and Hoosic River, a heavily forested area with strong interaction between produce fields and forest). Our data specifically suggests that in the watershed with widespread produce fields and sparse forest coverage, environmental selection, possibly caused by soil phosphorus, and slightly limited dispersal may result in relative heterogeneous composition of *E. coli* between produce field and forest sites and potential local adaptation in *E. coli*. In contrast, in the watershed with heavily forested areas, no evidence of environmental selection was observed, and dispersal facilitated by wildlife such as migratory bird flocks and small flocking insectivore/granivores may enhance the likelihood of genetic exchange among *E. coli* populations, resulting in relative homogeneous composition between produce field sites and forest sites in this watershed. This higher level of homogeneity is consistent with greater interaction between produce fields and forests in the Hoosic River watershed (adjacency_produce|forest_ = 36%) compared to Flint Creek (adjacency_produce|forest_ = 23%).

### Agricultural practice involving input of phosphorous in soil may enhance the selective pressure on *E. coli*

Agricultural activities normally involve cultivation and soil amendments, which could dramatically change soil organic matter and nutrient pools in comparison to undisturbed systems (e.g., forest) (42). Consequently, long-term organic and chemical amendments could dramatically impact the abundance, diversity, and composition of bacterial communities in soil of agricultural land (43). This is because such alteration of soil properties could trigger selective pressures on bacteria, sorting the individuals or traits that better cope with modified soil condition, which has been termed “local adaptation” (23). For example, copper‐amendment in agricultural soil has been found to significantly increase the frequency of copper‐resistant Gram-negative bacteria (44). Based on the results of our study reported here, agricultural practices may have caused selective pressure on soil *E. coli*, partially resulting in the distinct *E. coli* composition between produce fields sites and forest sites in the Flint Creek watershed. Consistent with our findings, Dusek et al. (22) observed *E. coli* population structures that differed between cropland and forest, with much lower prevalence of *E. coli* in cropland than forest. The diverse lifestyles and phenotypes of *E. coli* strains were thought to be caused by population expansion paired with differential niche adaptation under specific selective pressures in the last 5 million years (39). Our findings suggest that agriculture-stimulated selective pressures may contribute to *E. coli* diversification.

Besides directly yielding selective pressures on *E. coli*, soil property alterations caused by agricultural activities may also indirectly impact the adaptation of *E. coli* by changing the interaction with other microbial taxa in the community. A wealth of studies have shown that anthropogenic activities in agricultural land greatly change biological soil characteristics including the diversity, structure, and metabolic guilds of microbial communities (45–48). For example, *E. coli* has been reported to exhibit bacteriocin-mediated competitive interactions (49) and cooperative interactions using cross-feeding metabolic products with other taxa in the microbial community (50). Changes in microbial community structure and composition caused by agricultural activities thus possibly generate selective pressures on *E. coli* through altering the competition and cooperation behaviors with other microbial taxa.

The high correlation between phosphorous and antimony and the dissimilarity of *E. coli* composition observed in the watershed with widespread produce fields (Flint Creek) suggest that the alteration in these two soil variables may change the structure of *E. coli* populations in Flint Creek. Phosphorus is one of the soil variables well documented to dramatically change after the conversion of undisturbed systems to agriculture (42, 51). The input of phosphorus in fertilizer and manure to agricultural systems have been reported to often exceed the output in harvested crops (52). Phosphorus is a critical nutrient for the growth of bacteria and is part of many biomolecules in bacterial cells (e.g., DNA, phospholipids, polyphosphates, and ATP). Phosphorus availability could act as an important selective force driving divergence among bacterial populations. For example, Coleman et al. (21) identified a number of genes encoding functions related to phosphorus acquisition and metabolism (e.g., alkaline phosphatase, a pathway for phosphonate utilization, upregulation during phosphorus-starvation conditions) as significantly enriched in *Prochlorococcus* populations in locations with lower phosphate concentrations in North Atlantic and North Pacific subtropical gyres. Antimony is a toxic metalloid present widely at trace concentrations in natural soil (53, 54). Its concentration could be elevated or even reach contamination threshold in agricultural lands due to human activities (55). For example, application of lead arsenate pesticides in produce field can increase antimony concentration, since antimony is present as a contaminant in the antimony- and arsenic-containing ores used for pesticide manufacturing (56). A previous study has shown that increased antimony could prevent the growth of *E. coli, Bacillus subtilis* and *Staphylococcus aureus*, and may affect nitrogen cycle in soil by changing urease activity under neutral pH (57). However, the concentration of antimony detected in this studied was relatively low across all sites, and for some sites the concentration was below detection limit, thus the presence of antimony may not necessarily inhibit the growth of *E. coli* in soil studied here. Therefore, it is most likely that phosphorus represents agriculture-related selective pressures on *E. coli,* though other physical soil parameters may have also played a role. In addition, the fact that this correlation was overserved in Flint Creek but not in Hoosic River suggests that *E. coli* populations found in different watersheds and environments may differ in their adaptive traits (e.g., those associated with phosphorous).

### *E. coli* in a watershed with high forest coverage may experience very weak selective pressure and a proximity effect of forest

In this study, environmental selection tended to be very weak on *E. coli* in watershed with higher forest coverage (Hoosic River). This relatively week environmental selection might be because, compared to produce fields, plant cover and shading in forest could moderate perturbations in soil moisture, nutrients and temperature, thus providing more favorable and stable conditions with fewer environmental stressors for *E. coli* (4, 22). As previously proposed (4), soil in undisturbed temperate forests could act as potential habitat for long-period persistent, even resident *E. coli* populations rather than acting as a transient habitat. Albeit *E. coli* may be exposed to fewer or less intense stressors in undisturbed environments, as compared to disturbed ones, some factors such as temperature, moisture and nutrients have been shown to be correlated with *E. coli* density in forest (58, 59). Due to lack of niche differentiation caused by environmental selection, we observed more homogeneous *E. coli* compositions between forest and produce field in watershed with higher forest coverage. We also observed that *E. coli* was much more prevalent in the watershed with higher forest coverage (72%) as compared to the watershed with lower forest coverage (35%), consistent with previous findings by Dusek et al. (22).

The higher prevalence of *E. coli* in the watershed with higher forest coverage might be caused by proximity effect, which proposes that the likelihood of *E. coli* isolation from surrounding sites such as produce field increases with the proximity to forests (22). Such a proximity effect is formed by the spread of *E. coli* out of forests into surrounding areas, given that forest is a vital sink for *E. coli* (4). In addition, the large adjacency between forest and produce fields in the watershed with higher forest coverage, which indicates strong direct interactions between the two land covers, may enhance the proximity effect. Consistent with our findings, Dusek et al. (22) reported that *E. coli* was more prevalent in a landscape with greater forest coverage; this study specifically showed that *E. coli* was most prevalent in soils sampled in close proximity (0 to 38 m) of forests, but was up to 90% less prevalent when forest cover in the 250m radius was less than 7%. In addition to *E. coli*, such proximity effects of forest have also been reported for *Listeria monocytogenes* and other *Listeria* species. Weller et al. (60) found that with a 100m increase in the distance of a sampling site from forests, the likelihood of *L. monocytogenes* and other *Listeria* species isolation in croplands decreased by 14% and 16%, respectively.

### Watershed landscape could constrain or facilitate the dispersal of soil *E. coli* by influencing the movement of wildlife host

Wildlife, which is thought to be an important vehicle for the dispersal of foodborne pathogens between hosts and locations (36), could enable bacteria to overcome landscape barriers and make the dispersal of bacteria more active. Several studies have specifically indicated that wildlife is a major source of *E. coli* in surface waters and may contribute to the contamination of *E. coli* in rural watersheds and produce fields by defecation (11, 36, 61, 62). Landscape connectivity (i.e., the degree to which a landscape facilitates or prevents movement of organisms among resource patches) and particular landscape elements such as the structure of habitat (e.g., riparian corridor, terrestrial land, waterbody) have also previously been shown to influence dispersal of pathogens (63, 64) and the movement of wildlife (41). Wildlife-dependent dispersal of *E. coli* would thus be indirectly impacted by landscape. Our results showed that the dispersal of soil *E. coli* in the watershed with widespread produce fields was slightly limited, while the dispersal of soil *E. coli* in the watershed with high forest coverage tended to be largely facilitated by wildlife, highlighting the important role of watershed landscapes on the dispersal of *E. coli*. These influences may consequently shape the spatial patterns of pathogenic *E. coli* persistence and incidence (65). Consistent with our findings, Mechai et al. (66) showed evidence of the impact of landscape connectivity on the dispersal patterns of *Borrelia burgdorferi*, particularly rodent-associated strains, which is relevant to the spread of Lyme disease risk across locations.

Based on the above notions, our observation that the dispersal of soil *E. coli* in the watershed with widespread produce fields was relatively limited could be explained by a combination of environmental constraints (i.e., relatively strong environmental selection) in soil and poor connectivity of agricultural areas, which may impede the movement of wildlife that disperses *E. coli* (41). By contrast, the dispersal of soil *E. coli* in the watershed with high forest coverage tended to be promoted by wildlife. This may be because forest exhibits better connectivity and could provide passage and support to the movement of wildlife vehicles of *E. coli*. Besides wildlife, it is also possible that the dispersal of *E. coli* was directly influenced by the landscape elements of the two watersheds. Forest and most produce fields in Hoosic River, which is heavily forested, were both located in a floodplain. By contrast, forest in Flint Creek was in a floodplain but produce fields were not. Since during periods of high discharge, a floodplain normally experiences flooding, such events may facilitate the dispersal of *E. coli* between forest and produce field particularly in the Hoosic River watershed. This hypothesis is supported by a number of modelling studies, showing that the peak fecal bacteria levels during flooding can be more than 20 or even 50 times higher than prior to flooding (67–69). Future studies on comparing the distribution of *E. coli* before, during, and after flooding and assessing the correlation between flooding-associated landscape factors (e.g., elevation and patchiness) and distribution of *E. coli* are needed for an improved understanding of the impact of landscape on the microbial biogeography.

Migratory bird flocks, small flocking insectivore/granivores, and large nuisance wildlife were identified, in our study here, as potential vehicles that disperse *E. coli* and affect the distribution of *E. coli* in the watershed with high forest coverage. Migratory bird flocks (e.g., Canada goose) tend to have low movement cost in all land-use types except for heavy urban development areas (70). Small flocking insectivore/granivores (e.g., European starling) tend to have low movement cost in all land-use types except for open water and heavy urban development areas (70). Large nuisance wildlife (e.g., white tailed deer or feral swine) tend to have low movement cost in forests, scrublands, grasslands, wooded wetlands and cultivated croplands, while they can have high movement cost to cross roads (71). Each of these three classes of wildlife have been previously reported (34, 35, 72) to serve as dispersal vehicles of *E. coli* and have been considered public health concerns in terms of agricultural contamination. For example, a study of *E. coli* isolated from fecal samples of Canada geese over a year in Colorado, US reported that the prevalence for *E. coli* ranged from 2% during the coldest months to 94% during the warmest time of the year (72). European starlings, which are considered an invasive species in the United States and a nuisance pest to agriculture, were proposed to be a potential suitable reservoir and vector of *E. coli* O157:H7, and can carry and disseminate this human pathogen to cattle (34). In addition, deer feces were reported to contaminate fresh strawberries, being responsible for an outbreak of *E. coli* O157:H7 infections in Oregon (35). Forest is a relative stable environment with less disturbance of anthropologic activities, thus serving as an ideal living habitat for wild animals (73). Forest may provide easy transport pathways for small flocking insectivore/granivores and large nuisance wildlife to move around and support high density of migratory birds, which increases the chance of colonized *E. coli* dispersed from forest to adjacent produce fields.

### Conclusion

By comparing the biogeographic patterns of *E. coli* isolated from two watershed with distinct landscape characteristics in New York state, we showed that terrestrial landscape could impact the distribution of *E. coli* by adjusting the importance of environmental selection and dispersal. Environmental stress was identified as a possible strong contributor to local adaptation of *E. coli* in the watershed with widespread produce fields. On the other hand, wildlife-driven dispersal, which could facilitate genetic exchange, was identified as a major force in shaping *E. coli* populations in the watershed with high forest coverage. As such, our findings not only highlight the critical role of landscape in driving the biogeographic pattern of *E. coli* in perspective of ecology, but also open the possibility that the evolutionary forces (e.g., positive selection, genetic drift, gene flow) driving the diversification of *E. coli* vary by watershed landscape as well. In addition, our study suggests that due to the less intense environmental stress, frequent wildlife-facilitated dispersal, and the proximity effect of forest on *E. coli*, produce fields in watershed with high forest coverage may have higher risk in *E. coli* contamination. This information can inform spatial modeling of food contamination risk associated with produce fields in different watersheds, which can be used to modify pre-harvest product sampling strategies and produce harvest methods to account for the spatial structure in contamination risk in a produce field. Despite the contributions to the field of microbial ecology, our study has some non-negligible limitations. First, we did not differentiate commensal and pathogenic *E. coli*; however, biogeographic patterns and the impact of landscape may differ between the two groups. Second, we only assessed the coverage of forest and produce field as one landscape attribute using a relatively small set of sampling sites, while many other landscape attributes such as the size, diversity, and richness of patch size may also be important to the biogeographic pattern and adaptation of *E. coli*. Third, wildlife population/community structure, which could be strongly affected by land-use features (74), was not included in the competing models. Future studies warrant more intensive sampling efforts, sequencing techniques with higher discriminatory power (e.g., whole genome sequencing), and comprehensive assessment of a wider range of high order landscape attributes and wildlife characteristics to better understand the impact of the landscape on the biographic pattern and adaptation of *E. coli*. Such methodology development, ideally validated by experimental data, could considerably improve prediction of produce contamination risk based on the potential influence of landscape on the dispersal of *E. coli* to produce field, benefit the development of trade-off risk assessments of food contamination, and eventually help to decrease human exposure to pathogenic enteric bacteria.

## MATERIALS AND METHODS

### Study sites and soil collection

Two watersheds with different landscape patterns, Flint Creek and Hoosic River, located in the New York state, were selected for this study based on topography and land-cover composition. Flint Creek is an area with widespread vegetable and livestock production that is sparsely forested (69% produce field, 12% forest by area), whereas the Hoosic River watershed is a heavily forested area with interspersed produce production (28% produce field, 38% forest by area). Soil sampling was carried out between September 4 and October 10, 2012 on 7 farms comprising 19 produce field sites and in 16 forest sites (Fig. 1). For produce fields, two parallel 200 m transects were laid in each field, perpendicular to the forest boundary. Along each transect, five soil samples (at approximately 5 cm-depth) were collected at 50 m intervals using sterile scoops (Fisher Scientific, Hampton, NH) and sterile Whirl-Pak bags (Nasco, Fort Atkinson, WI). Latex gloves and disposable plastic boot covers (Nasco, Fort Atkinson, WI) were worn for sample collection. Gloves and boot covers were changed between each site, and gloves were disinfected with 70% ethanol prior to sample collection. A total of 278 soil samples were collected with 142 and 136 samples collected from the Flint Creek and the Hoosic River watershed, respectively. All samples were transported to the Food Safety Lab at Cornell University in an icebox. Samples were stored at 4 ± 2°C in dark and processed within 24 h of collection.

### Isolation of *E. coli*

*E. coli* were isolated from soil samples as previously described (23). Briefly, 8 g sieved soil was diluted 1:10 in EC medium with 4-methylumbelliferyl-β,D-glucuronide broth (EC-MUG). To maximize genetic diversity among recovered *E. coli* isolates, the suspension was subdivided among four 96-well microtiter plates for a total of 384 subsamples of approximately 180 uL each. Microtiter plates were incubated at 37°C. Bacteria from fluorescent wells were isolated on EC-MUG agar plates and were further tested with a standard biochemical assay for glutamate decarboxylase and beta-glucuronidase activity. Isolates that were positive for these two tests were presumptively identified as *E. coli*, which were confirmed by subsequent gene sequencing as detailed below. No *E. coli* isolates were detected in samples from 3 produce sites (Field 7, Field 9, and Field 10), and 2 forest sites (Forest F4 and Forest F18).

### DNA extraction, MLST genotyping and clonal groups

Genomic DNA was extracted from *E. coli* by alkaline lysis of biomass in 50 mM NaOH at 95°C. Two genes (*mdh* and *uidA*) were sequenced first from all isolates. Then, only the unique two-gene sequence types from each sample were subjected to additional sequencing in five additional genes (*aspC, clpX, icd, lysP, fadD*) by Sanger sequencing, performed by the Cornell University Life Sciences Core Laboratories Center. Evaluation of sequence read quality and assembly of forward and reverse reads were performed using Perl scripts, which iterated runs of phred and CAP3, respectively. Sequences with a probability of error of > 0.005 (Q score < 23) in terms of read quality were edited manually, where possible, or discarded. Assembled sequences of each MLST locus were aligned and trimmed to standard base positions matching the *E. coli* K-12 sequence type from the STEC Center website (http://www.shigatox.net) (23). Alignments of assembled sequences for isolates from Flint Creek and Hoosic River are available on GitHub (https://github.com/pbergholz/Dispersal-cost-modeling). The clonal groups of *E. coli* strains for Flint Creek and Hoosic River were determined based on MLST sequence types using the goeBURST analysis program (75) at single locus variant level.

### Remotely sensed data and soil property data

GPS coordinates of sites were imported into the Geographical Resources Analysis Support System (GRASS) geographic information system (GIS) environment. Map layers for land cover (National Land Cover Database [NLCD], 2006) and the digital elevation model (DEM; Shuttle Radar Topography Mission, 1-arc-second data set) were acquired from the U.S. Geological Survey (USGS) Earth Explorer geographical data bank (http://earthexplorer.usgs.gov/). Map layers for soil characteristics were acquired from the U.S. Department of Agriculture Soil Survey Geographic (SSURGO) database (http://soils.usda.gov/survey/geography/ssurgo/). Road and hydrologic line graphs were obtained from the Cornell University Geospatial Information Repository (CUGIR; http://cugir.mannlib.cornell.edu//).

Percent landcover and adjacency were estimated by using FRAGSTATS v. 3.3 to analyze landcover within a 2 km buffer surrounding the Flint Creek and Hoosic River, respectively (76). All land NLCD maps identified as pasturage was reclassified as cropland for our analyses. Percent adjacency was calculated as the proportion pixels in the NLCD map that were adjacent forest and produce field, compared to the total of non-self-adjacencies in 2 km buffer surrounding the waterway. For example, adjacency_produce|forest_ = 10% would indicate that 10% of the edges of produce fields abutted forest in a given area.

Organic matter, moisture, pH, aluminium, arsenic, boron, barium, calcium, cadmium, cobalt, chromium, copper, iron, potassium, magnesium, manganese, molybdenum, sodium, nickel, phosphorus, lead, sulphur, strontium, and zinc content of soil samples were measured at Cornell Nutrient Analysis Lab.

### Distribution of *E. coli* clonal groups and its relationship with geographic location

The Mann-Whitney test was used to determine if number of clonal groups differed significantly between soil samples from produce field sites and forest sites for Flint Creek and Hoosic River. Principal coordinate analysis (PCoA) was conducted using phyloseq package in R 3.6.0 to visualize the distribution of *E. coli* clonal groups among sites, based on Bray-Curtis distance. The 95% confidence ellipse in the PCoA plot assumes a multivariate normal distribution. Permutational multivariate analysis of variance (PERMANOVA) (77) was employed using the adonis function in R 3.6.0’s vegan package to test whether the centroids and dispersion of sample groups as defined by land-use (produce field or forest) are equivalent for both groups based on Bray-Curtis distance of *E. coli* clonal groups. PERMANOVA test statistic (F) and *p*-value were obtained by 9,999 permutations. Analysis of similarities (ANOSIM) (77) was employed using the anosim function in R 3.6.0’s vegan package to test whether there is a significant difference between two groups (produce field sties and forest sites) of sampling units based on the Bray-Curtis distance of *E. coli* clonal groups. ANOSIM test statistic (R) and p-value were obtained by 9,999 permutations. To test the sampling bias which possibly caused the large variation of *E. coli* clonal groups observed across sampling sites within each watershed, sites with a number of clonal groups ≤ 3 (Field 6, Field 8, and Forest F9) were excluded and this subset of samples were repeated for PCoA, PERMANOVA, and ANOSIM analyses.

Mantel tests were performed using vegan package in R 3.6.0 to assess the relationship between the biological dissimilarity of *E. coli* and geographic distance (9,999 permutations). Biological dissimilarity of *E. coli* clonal groups was calculated in Bray-Curtis distance using vegan package in R 3.6.0. Geographic distance between isolates was calculated from latitude and longitude, using the geopy module in Python 3.6.8. Linear regression analysis of biological dissimilarity of *E. coli* and geographic distance was performed in R 3.6.0. Distance-decay relationship was inferred from the slope and R^2^ of the linear regression. A steeper slope with a larger R^2^ value suggests stronger distance-decay relationship.

### The relationship between *E. coli* clonal groups and soil variables

Due to a lack of mineral soil to measure soil property after combusting away the organic matter, Forest F6, Forest F11, and Forest F16 were not included in analyses on the relationship between *E. coli* clonal groups and soil variables. After screening for covariation, soil variables with low levels of covariation (r < 0.7 and *p* < 0.05 in Pearson’s correlation analysis) were selected for variation partitioning analysis (VPA) using vegan package in R 3.6.0 to quantify the relative contribution of the environment effect and the geographical effect on the dissimilarity of *E. coli* clonal groups based on Bray-Curtis distance (78). Principal coordinates of neighbor matrices (PCNM) were used to represent spatial patterns based on GPS coordinates (79). By using ‘ordistep’ function in vegan package, a subset of PCNM variables which significantly explained variation in the dissimilarity of *E. coli* clonal groups was included in VPA. Four components of variations were calculated in VPA: i) pure contribution of environmental effect (R^2^_A_ – R^2^_G_); ii) pure geographical effect (R^2^_A_ – R^2^_E_); iii) spatially structured environmental effect (R^2^_G_ + R^2^_E_ – R^2^_A_); and iv) unexplained effect (1- R^2^_A_). R^2^_A_, R^2^_G_, and R^2^_E_ represent variation of the dissimilarity of *E. coli* clonal groups explained by all variables, spatial variables, and environmental variables, respectively.

Partial Mantel test was performed to examine the correlation between environmental dissimilarity of each soil variable and the dissimilarity of *E. coli* clonal groups independent of geographical influence using vegan package in R 3.6.0 (9,999 permutations). Dissimilarity of *E. coli* clonal groups was calculated in Bray-Curtis distance, and environmental dissimilarity was calculated in Euclidian distance. Soil variables with a *p* value < 0.05 in partial Mantel tests were defined as key soil variables. Mann-Whitney tests were further performed to determine if key soil variables differed significantly between soil samples from produce field sites and forest sites.

### Dispersal model formulation and selection

To predict the dispersal of *E. coli* across watershed landscapes, multiple dispersal models were developed to describe landscape effects by integrating remotely sensed and field-collected data into resistance surfaces for wildlife vehicles. Four common classes of wildlife vehicles including (i) large nuisance wildlife species, (ii) small mammals, (iii) small flocking insectivore/granivores, and (iv) migratory bird flocks were selected in this study.

Predicted dispersal among sites was calculated according to the equation below:

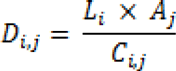

Where D*_i,j_* is the dispersal rate among sites *i* and *j*, L*_i_* is the *E. coli* load from the source site (i.e., starting point), A*j* is the attraction (gravity) coefficient of the sink site (i.e., stopping point) to vehicle and C*_i,j_* is the least-cost distance between sites *i* and *j*. *E. coli* load L*i* expresses the expected mobility of *E. coli* from these areas as a function of expected prevalence. Expected prevalence was inferred from random forest analysis of *E. coli* prevalence based on sampling excursions. One load map was generated per watershed. The attraction (gravity) coefficient A*j* describes the tendency of a dispersal vehicle to move towards an area on the landscape and expected residence-time of dispersal vehicle after they arrive at a location. Attraction was primarily a function of percent favored land-cover for each of the vehicles and interspersion of land-cover types. The least-cost distance C*_i,j_* describes the movement preferences of a dispersal vehicle in terms of a friction surface (borrowed from circuit theory) that predicts resistance of the landscape to movement of dispersal vehicles. The cost surfaces were a function of baseline resistance (dependent on the dispersal vehicle), riparian corridor effect (i.e., the tendency of wildlife to prefer movement through riparian forests), dispersal barrier effect (i.e., the strength of barriers to movement, such as major road- and water-ways), and proximity effect (i.e., the strength and type of edge interactions among forests, produce fields, pasturage, and urban areas). The least-cost distance was measured as the distance along the path that accrued the least cumulative cost between pairs of movement start and stop sites. The characteristics of the dispersal vehicles, *E. coli* load model (L_*i*_), attraction model (coefficient A_*j*_), cost model (i.e., riparian corridor effect, dispersal barrier effect and proximity effect) were shown in Table 1, which were summarized on the basis of published literature (73–84). Based on these characteristics, the most-likely attraction model and cost model were selected for each class of vehicle, generating the predicted dispersal models (Table 2).

For each of the predicted dispersal models for the four classes of wildlife vehicles, an association matrix D_*i,j*_ containing predicted dispersal rates along least cost paths among all pairs of sites was generated. This was accomplished by using a set of scripts developed in the GRASS GIS ver. 6.4.3 programming environment; Perl scripts were used to automate calculations in the GIS; scripts are available on GitHub (https://github.com/pbergholz/Dispersal-cost-modeling). Mantel tests were employed to estimate the correlation between predicted dispersal models and biological dissimilarity of *E. coli* clonal groups among sampled sites in each watershed using R version 3.6.0. Statistical significance of model fits was estimated by 9,999 permutations. The wildlife vehicle for which the predicted model had the highest significant correlation coefficient was deemed to represent the dominant dispersal vehicle for *E. coli*.

## Acknowledgements

This research was supported by the Center for Produce Safety (research agreement number 201121642-01, representing a subcontract under Award Number SCB11072 from the California Department of Food and Agriculture).

