## Supplemental material for "Adjacent terrestrial landscapes impact the biogeographical pattern of soil *Escherichia coli* in produce fields by modifying the importance of environmental selection and dispersal"

### Supplemental Materials

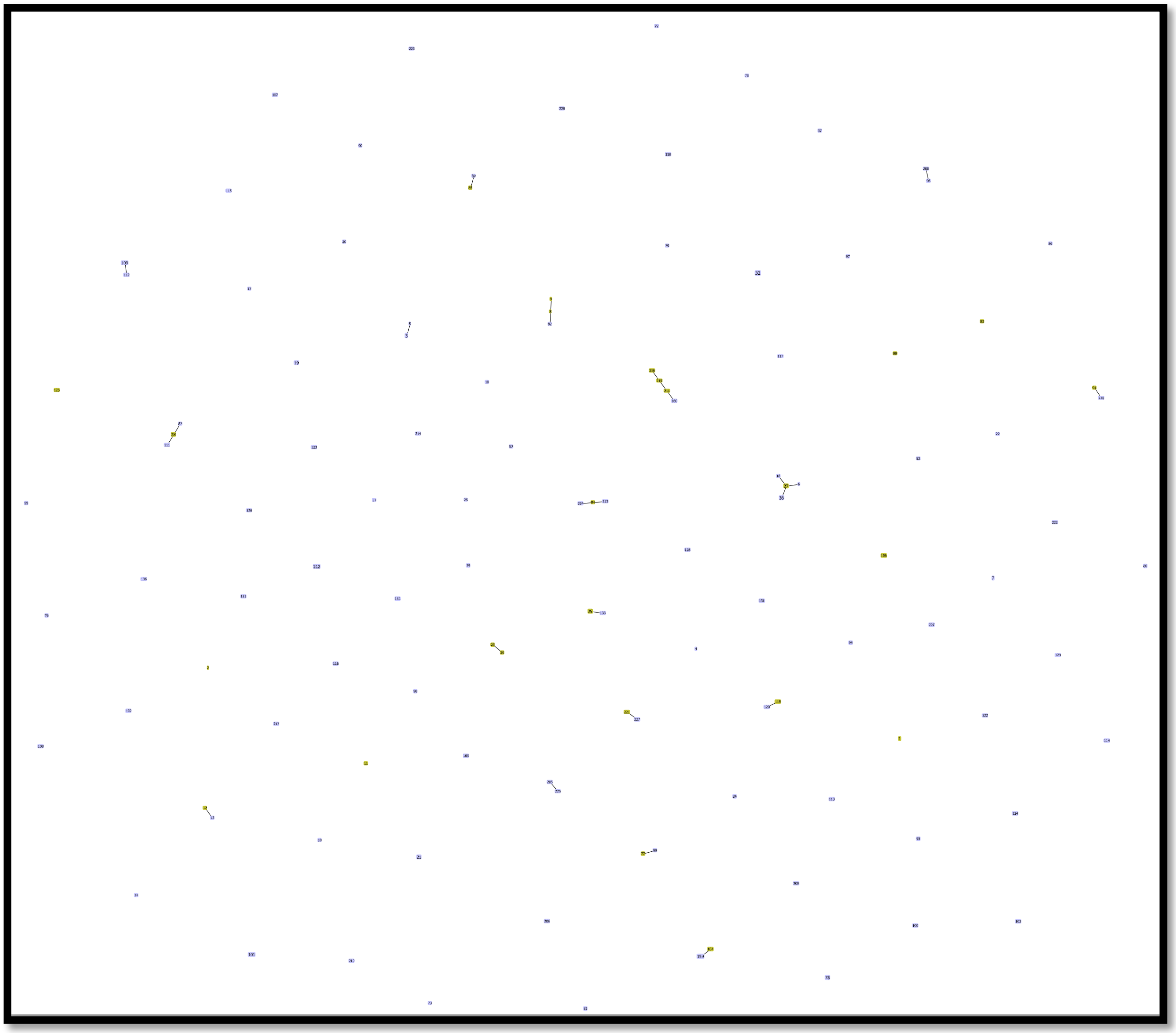

**FIG S1** The *E. coli* clonal groups in Flint Creek based on MLST sequence types at single locus variant level identified by goeBURST

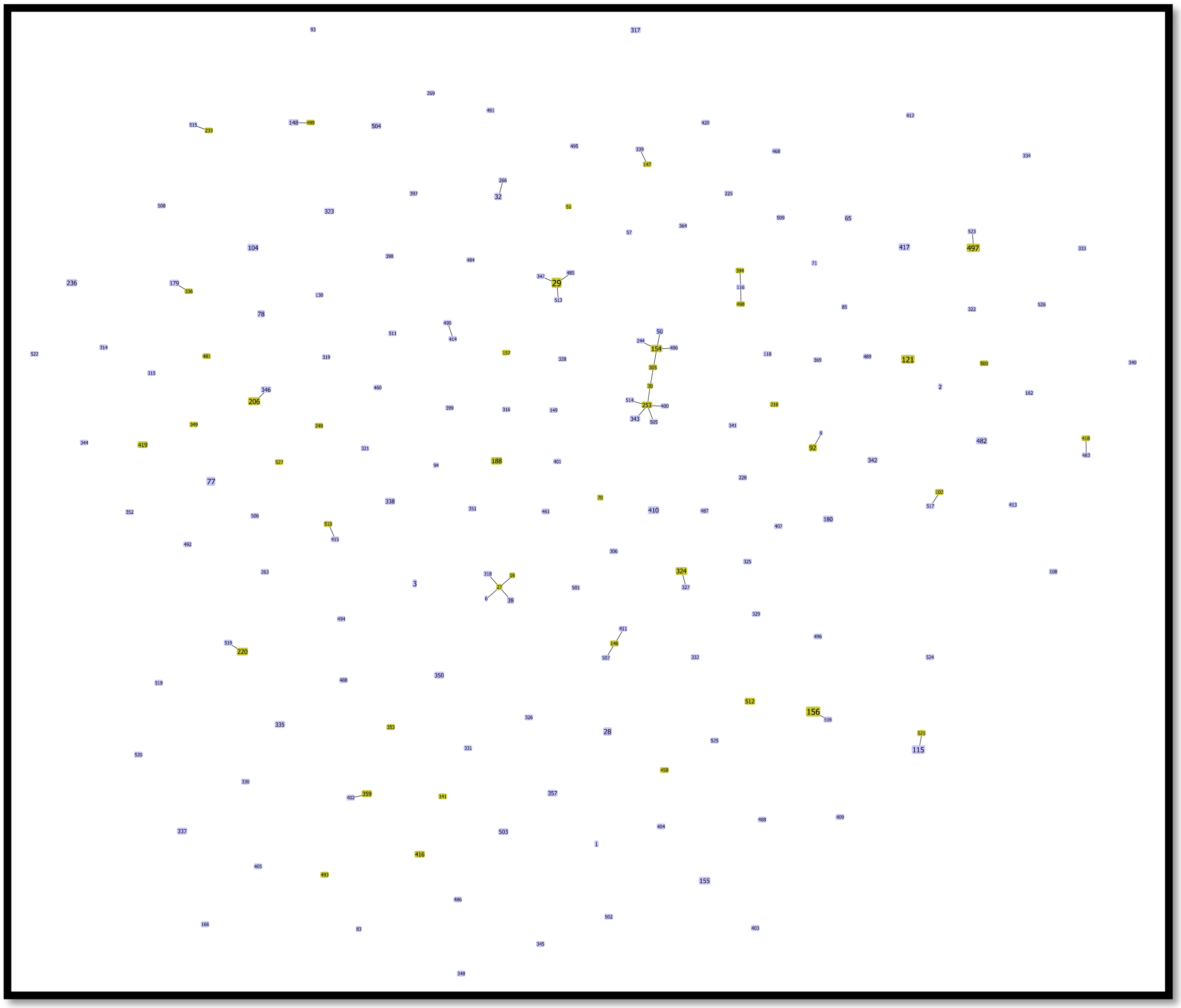

**FIG S2** The *E. coli* clonal groups in Hoosic River based on MLST sequence types at single locus variant level identified by goeBURST

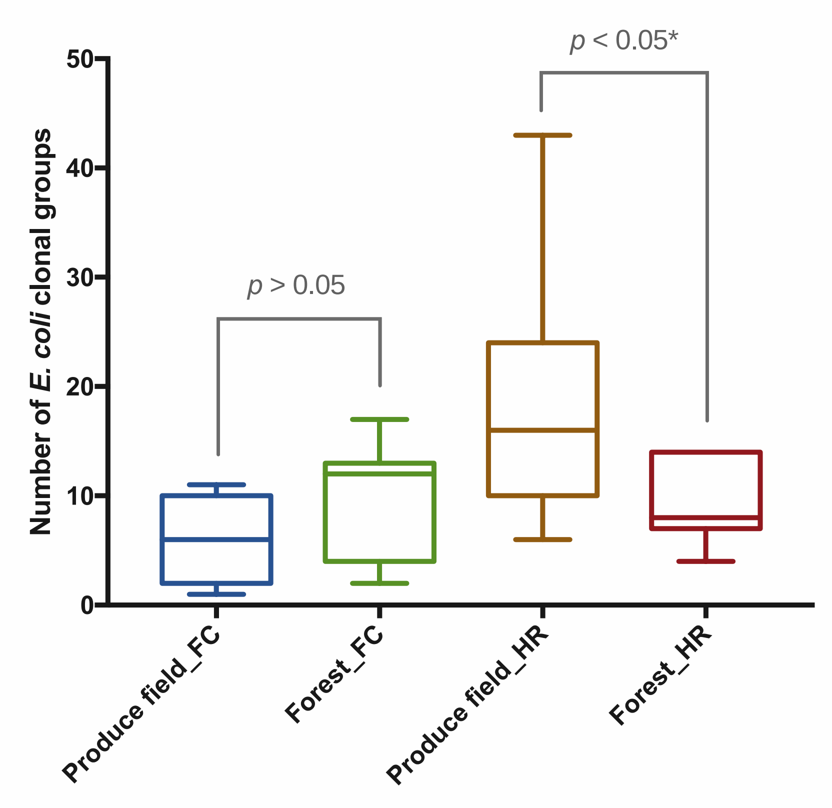


**FIG S3** Number of *E. coli* clonal groups in produce field sites and forest sites from Flint Creek (FC) and Hoosic River (HR). p values were determined by Mann-Whitney test. Minimum and maximum values are depicted by short horizontal lines above and below the box; the box signifies the upper and lower quartiles, and the mean is represented by a short line within the box.

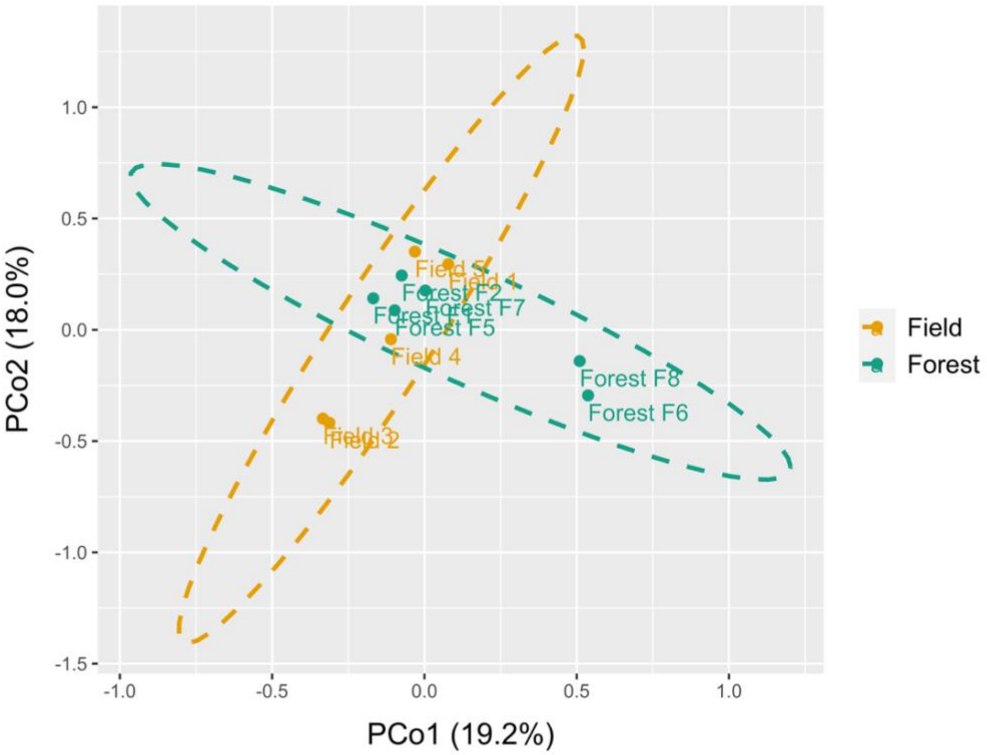


**FIG S4** PCoA plots of *E. coli* clonal groups of a subset of sites from Flint Creek. Field 6, Field 8, and Forest F9 were excluded in this analysis due to a low number of *E. coli* clonal groups detected (⩽ 3). Green dots indicated forest sites; orange dots indicated produce field sites; Green circle indicated the 95% confidence ellipse of forest sites; orange circle indicated the 95% confidence ellipse of produce field sites. PCo Axis 1 and 2 explained 19.2% and 18.0%, respectively, of the variation of *E. coli* clonal groups for Flint Creek.

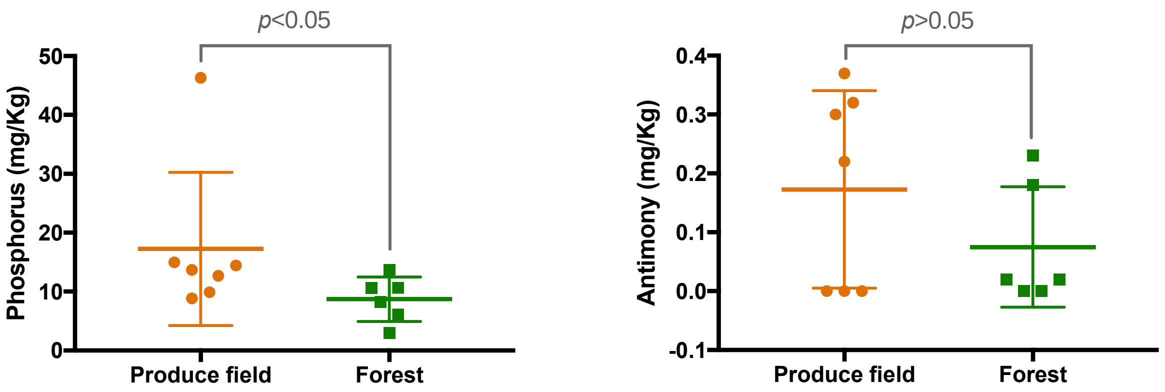


**FIG S5** (a) Phosphorus and (b) antimony of soil samples from produce field sites and forest sites of Flint Creek. *p* values were determined by Mann-Whitney test. Minimum and maximum values are depicted by horizontal line at the two ends of whiskers; points above and below the whiskers indicate outliers; and the mean is represented by a long horizontal line.

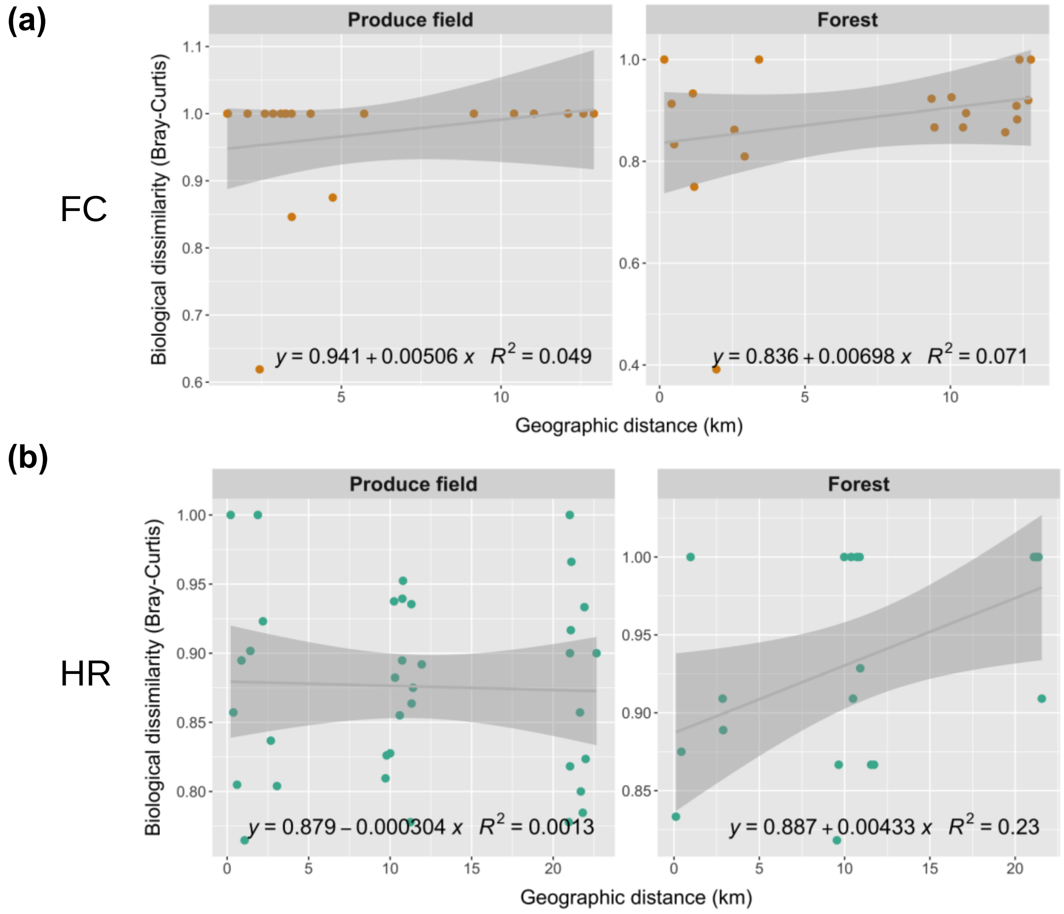


**FIG S6** Linear relationship between biological dissimilarity of *E. coli* clonal groups and geographical distance for produce field sites and forest sites in (a) Flint Creek (FC) and (b) Hoosic River (HR). The biological dissimilarity of *E. coli* clonal groups was calculated as Bray–Curtis distance. Geographical distance was calculated in actual physical distance. Linear regression line is in grey; shaded area indicates 95% confidence region; $R^2$ indicates the variability explained by fitted linear regression model and the formula of the linear relationship is shown.

**TABLE S1** Representative isolates of *E. coli* from unique MLST sequence types within each site in Flint Creek

| Isolate ID | Land use | Site ID | ST[a] |
|---|---|---|---|
| B71055 | Field | Field 1 | 205 |
| B71056 | Field | Field 1 | 101 |
| B71058 | Field | Field 1 | 206 |
| B71059 | Field | Field 1 | 109 |
| B71060 | Field | Field 1 | 207 |
| B71061 | Field | Field 1 | 208 |
| B71062 | Field | Field 1 | 159 |
| B71063 | Field | Field 1 | 28 |
| B71064 | Field | Field 1 | 3 |
| B71065 | Field | Field 1 | 29 |
| B71066 | Field | Field 1 | 160 |
| B71068 | Field | Field 1 | 209 |
| B71069 | Field | Field 1 | 210 |
| B71070 | Field | Field 1 | 211 |
| B71086 | Field | Field 2 | 217 |
| B71088 | Field | Field 2 | 219 |
| B71089 | Field | Field 2 | 21 |
| B71090 | Field | Field 2 | 220 |
| B71091 | Field | Field 2 | 221 |
| B71094 | Field | Field 2 | 222 |
| B71096 | Field | Field 2 | 223 |
| B71117 | Field | Field 3 | 57 |
| B71118 | Field | Field 3 | 226 |
| B71120 | Field | Field 3 | 136 |
| B71121 | Field | Field 3 | 227 |
| B70724 | Field | Field 4 | 32 |
| B70726 | Field | Field 4 | 128 |
| B70729 | Field | Field 4 | 129 |
| B70730 | Field | Field 4 | 130 |
| B70736 | Field | Field 4 | 131 |
| B70737 | Field | Field 4 | 132 |
| B70738 | Field | Field 4 | 133 |
| B70660 | Field | Field 5 | 94 |
| B70661 | Field | Field 5 | 95 |
| B70662 | Field | Field 5 | 96 |
| B70664 | Field | Field 5 | 97 |
| B70665 | Field | Field 5 | 98 |
| B70666 | Field | Field 5 | 99 |
| B70668 | Field | Field 5 | 100 |
| B70669 | Field | Field 5 | 101 |
| B70670 | Field | Field 5 | 102 |

| | | | |
|---|---|---|---|
| B70672 | Field | Field 5 | 103 |
| B70673 | Field | Field 5 | 104 |
| B70674 | Field | Field 5 | 105 |
| B70675 | Field | Field 5 | 32 |
| B70677 | Field | Field 5 | 106 |
| B70678 | Field | Field 5 | 107 |
| B70680 | Field | Field 5 | 108 |
| B70681 | Field | Field 5 | 27 |
| B70683 | Field | Field 5 | 109 |
| B70685 | Field | Field 5 | 111 |
| B70686 | Field | Field 5 | 112 |
| B70584 | Field | Field 6 | 72 |
| B70358 | Field | Field 8 | 1 |
| B70360 | Field | Field 8 | 2 |
| B71071 | Forest | Forest F1 | 3 |
| B71072 | Forest | Forest F1 | 159 |
| B71074 | Forest | Forest F1 | 212 |
| B71077 | Forest | Forest F1 | 213 |
| B71079 | Forest | Forest F1 | 214 |
| B71080 | Forest | Forest F1 | 215 |
| B71084 | Forest | Forest F1 | 216 |
| B71098 | Forest | Forest F2 | 212 |
| B71100 | Forest | Forest F2 | 224 |
| B71101 | Forest | Forest F2 | 155 |
| B71103 | Forest | Forest F2 | 225 |
| B71115 | Forest | Forest F2 | 38 |
| B70620 | Forest | Forest F5 | 84 |
| B70621 | Forest | Forest F5 | 85 |
| B70622 | Forest | Forest F5 | 37 |
| B70623 | Forest | Forest F5 | 86 |
| B70624 | Forest | Forest F5 | 87 |
| B70625 | Forest | Forest F5 | 88 |
| B70626 | Forest | Forest F5 | 38 |
| B70628 | Forest | Forest F5 | 89 |
| B70632 | Forest | Forest F5 | 90 |
| B70640 | Forest | Forest F5 | 19 |
| B70645 | Forest | Forest F5 | 91 |
| B70651 | Forest | Forest F5 | 92 |
| B70653 | Forest | Forest F5 | 93 |
| B70689 | Forest | Forest F5 | 78 |
| B70690 | Forest | Forest F5 | 113 |
| B70693 | Forest | Forest F5 | 114 |
| B70695 | Forest | Forest F5 | 115 |
| B70696 | Forest | Forest F5 | 116 |

| | | | |
|---|---|---|---|
| B70697 | Forest | Forest F5 | 117 |
| B70698 | Forest | Forest F5 | 118 |
| B70699 | Forest | Forest F5 | 51 |
| B70700 | Forest | Forest F5 | 119 |
| B70705 | Forest | Forest F5 | 120 |
| B70709 | Forest | Forest F5 | 121 |
| B70713 | Forest | Forest F5 | 122 |
| B70714 | Forest | Forest F5 | 1 |
| B70716 | Forest | Forest F5 | 123 |
| B70717 | Forest | Forest F5 | 124 |
| B70718 | Forest | Forest F5 | 125 |
| B70719 | Forest | Forest F5 | 32 |
| B70721 | Forest | Forest F5 | 126 |
| B70723 | Forest | Forest F5 | 127 |
| B70585 | Forest | Forest F6 | 73 |
| B70586 | Forest | Forest F6 | 74 |
| B70587 | Forest | Forest F6 | 75 |
| B70589 | Forest | Forest F6 | 76 |
| B70592 | Forest | Forest F6 | 7 |
| B70593 | Forest | Forest F6 | 77 |
| B70594 | Forest | Forest F6 | 78 |
| B70600 | Forest | Forest F6 | 79 |
| B70601 | Forest | Forest F6 | 30 |
| B70603 | Forest | Forest F6 | 80 |
| B70610 | Forest | Forest F6 | 81 |
| B70613 | Forest | Forest F6 | 82 |
| B70614 | Forest | Forest F6 | 83 |
| B70384 | Forest | Forest F7 | 11 |
| B70385 | Forest | Forest F7 | 12 |
| B70388 | Forest | Forest F7 | 13 |
| B70389 | Forest | Forest F7 | 14 |
| B70395 | Forest | Forest F7 | 16 |
| B70398 | Forest | Forest F7 | 17 |
| B70399 | Forest | Forest F7 | 18 |
| B70400 | Forest | Forest F7 | 19 |
| B70402 | Forest | Forest F7 | 20 |
| B70404 | Forest | Forest F7 | 21 |
| B70406 | Forest | Forest F7 | 22 |
| B70408 | Forest | Forest F7 | 23 |
| B70409 | Forest | Forest F7 | 24 |
| B70411 | Forest | Forest F7 | 25 |
| B70416 | Forest | Forest F7 | 26 |
| B70418 | Forest | Forest F7 | 10 |
| B70419 | Forest | Forest F7 | 27 |

| | | | |
|---|---|---|---|
| B70420 | Forest | Forest F7 | 28 |
| B70421 | Forest | Forest F7 | 29 |
| B70361 | Forest | Forest F8 | 3 |
| B70362 | Forest | Forest F8 | 4 |
| B70363 | Forest | Forest F8 | 5 |
| B70364 | Forest | Forest F8 | 6 |
| B70366 | Forest | Forest F8 | 7 |
| B70371 | Forest | Forest F9 | 8 |
| B70376 | Forest | Forest F9 | 9 |

[a]MLST sequence type

**TABLE S2** Representative isolates of *E. coli* from unique MLST sequence types within each site in Hoosic River

| Isolate ID | Land use | Site ID | ST[a] |
|---|---|---|---|
| B71506 | Field | Field 11 | 28 |
| B71507 | Field | Field 11 | 38 |
| B71512 | Field | Field 11 | 306 |
| B71514 | Field | Field 11 | 77 |
| B71515 | Field | Field 11 | 121 |
| B71516 | Field | Field 11 | 216 |
| B72012 | Field | Field 12 | 1 |
| B72013 | Field | Field 12 | 397 |
| B72015 | Field | Field 12 | 78 |
| B72017 | Field | Field 12 | 6 |
| B72018 | Field | Field 12 | 29 |
| B72020 | Field | Field 12 | 398 |
| B72022 | Field | Field 12 | 154 |
| B72023 | Field | Field 12 | 399 |
| B71538 | Field | Field 13 | 314 |
| B71541 | Field | Field 13 | 315 |
| B71546 | Field | Field 13 | 316 |
| B71548 | Field | Field 13 | 317 |
| B71550 | Field | Field 13 | 318 |
| B71557 | Field | Field 13 | 319 |
| B71558 | Field | Field 13 | 249 |
| B71561 | Field | Field 13 | 321 |
| B71562 | Field | Field 13 | 322 |
| B71571 | Field | Field 13 | 236 |
| B71572 | Field | Field 13 | 225 |
| B71574 | Field | Field 13 | 156 |
| B71575 | Field | Field 13 | 323 |
| B71576 | Field | Field 13 | 324 |
| B71577 | Field | Field 13 | 325 |
| B71579 | Field | Field 13 | 326 |
| B71581 | Field | Field 13 | 236 |
| B71582 | Field | Field 13 | 327 |
| B71584 | Field | Field 13 | 28 |
| B71586 | Field | Field 13 | 29 |
| B71592 | Field | Field 13 | 328 |
| B71596 | Field | Field 13 | 329 |
| B71597 | Field | Field 13 | 121 |
| B71599 | Field | Field 13 | 78 |
| B71600 | Field | Field 13 | 324 |
| B71605 | Field | Field 13 | 323 |
| B71606 | Field | Field 13 | 104 |

| | | | |
|---|---|---|---|
| B71608 | Field | Field 13 | 330 |
| B71610 | Field | Field 13 | 331 |
| B71611 | Field | Field 13 | 317 |
| B71615 | Field | Field 13 | 332 |
| B71617 | Field | Field 13 | 333 |
| B71656 | Field | Field 13 | 343 |
| B71666 | Field | Field 13 | 77 |
| B71668 | Field | Field 13 | 206 |
| B71669 | Field | Field 13 | 157 |
| B71672 | Field | Field 13 | 116 |
| B71673 | Field | Field 13 | 343 |
| B71674 | Field | Field 13 | 115 |
| B71677 | Field | Field 13 | 344 |
| B71679 | Field | Field 13 | 155 |
| B71680 | Field | Field 13 | 104 |
| B71682 | Field | Field 13 | 345 |
| B71684 | Field | Field 13 | 346 |
| B71685 | Field | Field 13 | 347 |
| B71689 | Field | Field 13 | 1 |
| B71690 | Field | Field 13 | 78 |
| B71692 | Field | Field 13 | 324 |
| B71693 | Field | Field 13 | 156 |
| B71700 | Field | Field 13 | 348 |
| B71701 | Field | Field 13 | 188 |
| B71702 | Field | Field 13 | 349 |
| B71703 | Field | Field 13 | 244 |
| B71704 | Field | Field 13 | 29 |
| B71706 | Field | Field 13 | 350 |
| B71707 | Field | Field 13 | 121 |
| B71710 | Field | Field 13 | 93 |
| B71711 | Field | Field 13 | 351 |
| B71715 | Field | Field 13 | 352 |
| B72118 | Field | Field 14 | 337 |
| B72119 | Field | Field 14 | 206 |
| B72122 | Field | Field 14 | 179 |
| B72124 | Field | Field 14 | 65 |
| B72125 | Field | Field 14 | 188 |
| B72128 | Field | Field 14 | 141 |
| B72129 | Field | Field 14 | 108 |
| B72132 | Field | Field 14 | 38 |
| B72139 | Field | Field 14 | 415 |
| B72141 | Field | Field 14 | 416 |
| B72143 | Field | Field 14 | 29 |
| B72144 | Field | Field 14 | 417 |
| B72147 | Field | Field 14 | 51 |

| | | | |
|---|---|---|---|
| B72149 | Field | Field 14 | 77 |
| B72170 | Field | Field 14 | 416 |
| B72534 | Field | Field 15 | 353 |
| B72535 | Field | Field 15 | 180 |
| B72536 | Field | Field 15 | 147 |
| B72537 | Field | Field 15 | 481 |
| B72538 | Field | Field 15 | 359 |
| B72539 | Field | Field 15 | 2 |
| B72541 | Field | Field 15 | 482 |
| B72544 | Field | Field 15 | 483 |
| B72545 | Field | Field 15 | 484 |
| B72546 | Field | Field 15 | 208 |
| B72547 | Field | Field 15 | 180 |
| B72548 | Field | Field 15 | 485 |
| B72549 | Field | Field 15 | 206 |
| B72551 | Field | Field 15 | 482 |
| B72552 | Field | Field 15 | 2 |
| B72555 | Field | Field 15 | 154 |
| B72556 | Field | Field 15 | 486 |
| B72559 | Field | Field 15 | 77 |
| B72563 | Field | Field 15 | 410 |
| B72565 | Field | Field 15 | 71 |
| B72566 | Field | Field 15 | 104 |
| B72569 | Field | Field 15 | 487 |
| B72573 | Field | Field 15 | 410 |
| B72578 | Field | Field 15 | 417 |
| B72579 | Field | Field 15 | 488 |
| B72582 | Field | Field 15 | 346 |
| B72616 | Field | Field 15 | 495 |
| B72617 | Field | Field 15 | 220 |
| B72618 | Field | Field 15 | 70 |
| B72619 | Field | Field 15 | 208 |
| B72627 | Field | Field 15 | 417 |
| B72628 | Field | Field 15 | 8 |
| B72629 | Field | Field 15 | 496 |
| B72630 | Field | Field 16 | 263 |
| B72631 | Field | Field 16 | 29 |
| B72632 | Field | Field 16 | 148 |
| B72633 | Field | Field 16 | 497 |
| B72634 | Field | Field 16 | 65 |
| B72635 | Field | Field 16 | 156 |
| B72637 | Field | Field 16 | 77 |
| B72638 | Field | Field 16 | 29 |
| B72639 | Field | Field 16 | 29 |
| B72647 | Field | Field 16 | 149 |

| | | | |
|---|---|---|---|
| B72648 | Field | Field 16 | 498 |
| B72649 | Field | Field 16 | 148 |
| B72650 | Field | Field 16 | 121 |
| B72651 | Field | Field 16 | 499 |
| B72661 | Field | Field 16 | 500 |
| B72662 | Field | Field 16 | 166 |
| B72688 | Field | Field 17 | 369 |
| B72689 | Field | Field 17 | 503 |
| B72690 | Field | Field 17 | 504 |
| B72691 | Field | Field 17 | 338 |
| B72694 | Field | Field 17 | 503 |
| B72695 | Field | Field 17 | 85 |
| B72696 | Field | Field 17 | 253 |
| B72705 | Field | Field 17 | 505 |
| B72706 | Field | Field 17 | 146 |
| B72707 | Field | Field 17 | 506 |
| B72708 | Field | Field 17 | 507 |
| B72710 | Field | Field 17 | 508 |
| B72711 | Field | Field 17 | 509 |
| B72714 | Field | Field 17 | 305 |
| B72719 | Field | Field 17 | 342 |
| B72720 | Field | Field 17 | 3 |
| B72728 | Field | Field 17 | 458 |
| B72729 | Field | Field 17 | 419 |
| B72731 | Field | Field 17 | 228 |
| B72732 | Field | Field 17 | 510 |
| B72748 | Field | Field 18 | 461 |
| B72749 | Field | Field 18 | 156 |
| B72752 | Field | Field 18 | 115 |
| B72753 | Field | Field 18 | 497 |
| B72754 | Field | Field 18 | 513 |
| B72756 | Field | Field 18 | 514 |
| B72758 | Field | Field 18 | 115 |
| B72759 | Field | Field 18 | 118 |
| B72760 | Field | Field 18 | 497 |
| B72763 | Field | Field 18 | 27 |
| B72764 | Field | Field 18 | 156 |
| B72767 | Field | Field 18 | 121 |
| B72774 | Field | Field 18 | 102 |
| B72775 | Field | Field 18 | 515 |
| B72777 | Field | Field 18 | 516 |
| B72779 | Field | Field 18 | 517 |
| B72780 | Field | Field 18 | 518 |
| B72781 | Field | Field 18 | 220 |
| B72782 | Field | Field 18 | 519 |

| | | | |
|---|---|---|---|
| B72785 | Field | Field 18 | 520 |
| B72786 | Field | Field 18 | 512 |
| B72787 | Field | Field 18 | 504 |
| B72788 | Field | Field 18 | 115 |
| B72789 | Field | Field 18 | 521 |
| B72790 | Field | Field 18 | 522 |
| B72791 | Field | Field 18 | 523 |
| B72792 | Field | Field 18 | 524 |
| B72794 | Field | Field 18 | 497 |
| B72796 | Field | Field 18 | 525 |
| B72797 | Field | Field 18 | 526 |
| B72798 | Field | Field 18 | 527 |
| B72799 | Field | Field 18 | 156 |
| B72801 | Field | Field 18 | 220 |
| B72806 | Field | Field 18 | 497 |
| B72808 | Field | Field 18 | 236 |
| B72052 | Field | Field 19 | 253 |
| B72055 | Field | Field 19 | 406 |
| B72060 | Field | Field 19 | 3 |
| B72067 | Field | Field 19 | 335 |
| B72070 | Field | Field 19 | 154 |
| B72071 | Field | Field 19 | 92 |
| B72077 | Field | Field 19 | 179 |
| B72080 | Field | Field 19 | 407 |
| B72081 | Field | Field 19 | 92 |
| B72083 | Field | Field 19 | 408 |
| B72086 | Field | Field 19 | 30 |
| B72088 | Field | Field 19 | 409 |
| B72091 | Field | Field 19 | 156 |
| B72097 | Field | Field 19 | 156 |
| B72098 | Field | Field 19 | 92 |
| B72099 | Field | Field 19 | 357 |
| B72100 | Field | Field 19 | 410 |
| B72101 | Field | Field 19 | 32 |
| B72102 | Field | Field 19 | 411 |
| B72103 | Field | Field 19 | 364 |
| B72104 | Field | Field 19 | 412 |
| B72106 | Field | Field 19 | 413 |
| B72108 | Field | Field 19 | 414 |
| B72110 | Field | Field 19 | 155 |
| B71518 | Forest | Forest F11 | 307 |
| B71519 | Forest | Forest F11 | 92 |
| B71520 | Forest | Forest F11 | 308 |
| B71524 | Forest | Forest F11 | 309 |
| B71527 | Forest | Forest F11 | 253 |

| | | | |
|---|---|---|---|
| B71528 | Forest | Forest F11 | 38 |
| B71529 | Forest | Forest F11 | 236 |
| B71530 | Forest | Forest F11 | 159 |
| B71531 | Forest | Forest F11 | 30 |
| B71532 | Forest | Forest F11 | 160 |
| B71533 | Forest | Forest F11 | 310 |
| B71534 | Forest | Forest F11 | 311 |
| B71535 | Forest | Forest F11 | 312 |
| B71537 | Forest | Forest F11 | 313 |
| B72024 | Forest | Forest F12 | 400 |
| B72025 | Forest | Forest F12 | 401 |
| B72026 | Forest | Forest F12 | 402 |
| B72030 | Forest | Forest F12 | 3 |
| B72034 | Forest | Forest F12 | 403 |
| B72042 | Forest | Forest F12 | 269 |
| B72044 | Forest | Forest F12 | 404 |
| B72047 | Forest | Forest F12 | 405 |
| B71622 | Forest | Forest F13 | 29 |
| B71624 | Forest | Forest F13 | 334 |
| B71628 | Forest | Forest F13 | 335 |
| B71630 | Forest | Forest F13 | 28 |
| B71631 | Forest | Forest F13 | 336 |
| B71633 | Forest | Forest F13 | 337 |
| B71640 | Forest | Forest F13 | 338 |
| B71641 | Forest | Forest F13 | 50 |
| B71644 | Forest | Forest F13 | 339 |
| B71647 | Forest | Forest F13 | 340 |
| B71648 | Forest | Forest F13 | 341 |
| B71649 | Forest | Forest F13 | 94 |
| B71650 | Forest | Forest F13 | 121 |
| B71651 | Forest | Forest F13 | 50 |
| B71652 | Forest | Forest F13 | 342 |
| B72150 | Forest | Forest F14 | 418 |
| B72151 | Forest | Forest F14 | 266 |
| B72158 | Forest | Forest F14 | 419 |
| B72159 | Forest | Forest F14 | 394 |
| B72162 | Forest | Forest F14 | 3 |
| B72165 | Forest | Forest F14 | 16 |
| B72167 | Forest | Forest F14 | 420 |
| B72583 | Forest | Forest F15 | 489 |
| B72587 | Forest | Forest F15 | 29 |
| B72588 | Forest | Forest F15 | 490 |
| B72591 | Forest | Forest F15 | 491 |
| B72594 | Forest | Forest F15 | 492 |
| B72596 | Forest | Forest F15 | 155 |

| | | | |
|---|---|---|---|
| B72598 | Forest | Forest F15 | 493 |
| B72602 | Forest | Forest F15 | 206 |
| B72603 | Forest | Forest F15 | 357 |
| B72605 | Forest | Forest F15 | 162 |
| B72606 | Forest | Forest F15 | 460 |
| B72608 | Forest | Forest F15 | 494 |
| B72609 | Forest | Forest F15 | 29 |
| B72613 | Forest | Forest F15 | 468 |
| B72615 | Forest | Forest F15 | 359 |
| B72672 | Forest | Forest F16 | 32 |
| B72673 | Forest | Forest F16 | 501 |
| B72676 | Forest | Forest F16 | 115 |
| B72678 | Forest | Forest F16 | 502 |
| B72679 | Forest | Forest F16 | 77 |
| B72682 | Forest | Forest F16 | 83 |
| B72686 | Forest | Forest F16 | 350 |
| B72687 | Forest | Forest F16 | 188 |
| B72733 | Forest | Forest F17 | 57 |
| B72734 | Forest | Forest F17 | 32 |
| B72736 | Forest | Forest F17 | 482 |
| B72738 | Forest | Forest F17 | 511 |
| B72739 | Forest | Forest F17 | 28 |
| B72740 | Forest | Forest F17 | 233 |
| B72742 | Forest | Forest F17 | 130 |
| B72745 | Forest | Forest F17 | 512 |

[a]MLST sequence type

**TABLE S3** PERMANOVA of presence/absence of *E. coli* clonal groups in Flint Creek and Hoosic River [a]

| Watershed | Grouping factor[b] | F Statistic | *p*-value |
|---|---|---|---|
| **Flint Creek** | **Land use type** | **1.704** | **0.015** |
| Hoosic River | Land use type | 0.792 | 0.857 |

[a] Rows in boldface indicate that PERMANOVA test was significant ($p < 0.05$).
[b] Land use type – forest vs produce field.

**TABLE S4** ANOSIM of presence/absence of *E. coli* clonal groups in Flint Creek and Hoosic River [a]

| Watershed | Grouping factor[b] | R Statistic | *p*-value |
|---|---|---|---|
| **Flint Creek** | **Land use type** | **0.200** | **0.035** |
| Hoosic River | Land use type | -0.031 | 0.580 |

[a] Rows in boldface indicate that ANOSIM test was significant ($p < 0.05$).
[b] Land use type – forest vs produce field.

**TABLE S5** PERMANOVA and ANOSIM of presence/absence of *E. coli* clonal groups in Flint Creek based on a subset of samples [a]

| Test | Grouping factor[b] | Statistic | *p*-value |
|---|---|---|---|
| PERMANOVA | Land use type | 1.441 | 0.050 |
| ANOSIM | Land use type | 0.176 | 0.088 |

[a] Samples with a number of clonal groups ≤ 3 (Field 6, Field 8, and Forest F9) were excluded in this analysis.
[b] Land use type – forest vs produce field.

**TABLE S6** Soil variables selected for VPA and partial Mantel tests

| Site | Moisture (%) | pH | Sodium (mg/Kg) | Phosphorus (mg/Kg) | Barium (mg/Kg) | Manganese (mg/Kg) | Antimony (mg/Kg) |
|---|---|---|---|---|---|---|---|
| | | | | Flint Creek | | | |
| Field 1 | 2.60 | 5.95 | 514.83 | 13.67 | 21.52 | 7.65 | 0.00 |
| Field 2 | 0.83 | 6.73 | 278.60 | 8.87 | 16.81 | 5.95 | 0.37 |
| Field 3 | 0.67 | 7.14 | 350.69 | 9.88 | 12.54 | 5.69 | 0.30 |
| Field 4 | 2.14 | 7.28 | 523.95 | 46.33 | 10.27 | 5.98 | 0.00 |
| Field 5 | 0.71 | 6.40 | 373.19 | 14.42 | 11.94 | 8.01 | 0.00 |
| Field 6 | 0.49 | 7.53 | 245.40 | 12.69 | 15.17 | 8.18 | 0.22 |
| Field 8 | 0.62 | 6.83 | 613.95 | 14.98 | 11.29 | 8.75 | 0.32 |
| Forest F1 | 1.60 | 7.63 | 2891.90 | 10.63 | 21.29 | 21.37 | 0.00 |
| Forest F2 | 2.60 | 5.95 | 514.83 | 13.67 | 21.52 | 7.65 | 0.00 |
| Forest F5 | 0.92 | 6.67 | 249.93 | 6.12 | 14.01 | 14.85 | 0.02 |
| Forest F7 | 0.84 | 7.41 | 536.64 | 10.60 | 19.76 | 34.12 | 0.02 |
| Forest F8 | 0.85 | 6.82 | 294.99 | 2.99 | 10.20 | 10.92 | 0.23 |
| Forest F9 | 1.71 | 6.88 | 350.94 | 8.29 | 20.90 | 17.34 | 0.18 |
| | | | | Hoosic River | | | |
| Field 11 | 0.78 | 7.25 | 360.56 | 3.58 | 25.75 | 13.02 | 0.40 |
| Field 12 | 0.66 | 7.6 | 291.28 | 14.54 | 8.64 | 15.05 | 0.23 |
| Field 13 | 0.60 | 6.52 | 294.72 | 4.85 | 23.63 | 8.89 | 0.35 |
| Field 14 | 0.49 | 6.11 | 262.11 | 7.49 | 17.35 | 18.09 | 0.49 |
| Field 15 | 0.59 | 6.38 | 233.12 | 8.12 | 12.86 | 6.46 | 0.33 |
| Field 16 | 0.55 | 6.67 | 227.21 | 11.69 | 16.40 | 4.68 | 0.25 |
| Field 17 | 0.43 | 6.60 | 166.68 | 11.82 | 13.14 | 5.18 | 0.29 |
| Field 18 | 0.56 | 7.31 | 276.95 | 12.12 | 17.86 | 5.96 | 0.46 |
| Field 19 | 0.63 | 6.75 | 334.94 | 37.84 | 15.35 | 5.92 | 0.21 |
| Forest F12 | 0.52 | 7.32 | 233.20 | 5.09 | 19.73 | 10.55 | 0.42 |
| Forest F13 | 0.49 | 7.46 | 457.78 | 7.43 | 18.12 | 20.90 | 0.25 |
| Forest F14 | 0.64 | 7.30 | 398.22 | 14.67 | 19.62 | 20.58 | 0.09 |
| Forest F15 | 0.87 | 5.56 | 210.43 | 6.98 | 9.23 | 18.92 | 0.23 |
| Forest F17 | 0.47 | 7.41 | 230.53 | 3.39 | 16.02 | 13.44 | 0.34 |

**TABLE S7** Mantel correlations between the dissimilarity of *E. coli* clonal groups based on Bray-Curtis distance and geographic distance for sites in Flint Creek and Hoosic River

| Watershed | R[a] | *p*-value |
|---|---|---|
| Flint Creek | 0.163 | 0.080 |
| Hoosic River | 0.106 | 0.124 |

[a] R: Mantel correlation coefficient

**TABLE S8** Mantel correlations between the dissimilarity of *E. coli* clonal groups based on Bray-Curtis distance and geographic distance for produce field sites and forest sites in Flint Creek and Hoosic River[a]

| Land-use | R[b] | *p*-value |
|---|---|---|
| | Flint Creek | |
| Produce field | 0.222 | 0.132 |
| Forest | 0.267 | 0.134 |
| | Hoosic River | |
| Produce field | -0.04 | 0.558 |
| **Forest** | **0.485** | **0.016** |

[a] Rows in boldface indicate that Mantel test was significant at *p* level of 0.05.
[b] R: Mantel correlation coefficient
